# Unambiguous assignment of kinked beta sheets leads to insights into molecular grammar of reversibility in biomolecular condensates

**DOI:** 10.1101/2024.12.05.627008

**Authors:** Irawati Roy, Rajeswari Appadurai, Anand Srivastavava

## Abstract

Kinked-*β* sheets are short peptide motifs that appear as distortions in *β*-strands and often mediate formation of reversible amyloid fibrils in prion-like proteins. Standard methods for assigning secondary structures cannot distinguish these esoteric motifs. Here, we provide a supervised machine learning based structural quantification map to unambiguously characterize Kinked-*β* sheets from coordinate data. We find that these motifs, although deviating from standard *β*-strand region of the Ramachandran plot, scatter around the allowed regions. We also demonstrate the applicability of our technique in wresting out LARKS, which are kinked *β*-strands with designated sequence. Additionally, from our exhaustive simulation generated conformations, we create a repository of potential kinked peptide-segments that can be used as a screening-library for assigning beta-kinks in unresolved coordinate dataset. Overall, our map for Kinked-*β* provides a robust framework for detailed structural and kinetics investigation of these important motifs in prion-like proteins that lead to formation of amyloid fibrils.

## Introduction

Biomolecular condensates are membrane-less organelles (MLOs) that enable numerous complex functions by tightly regulating biochemical reactions in space and time in the crowded cellular milieu.^1^ Cytoplasmic Processing Bodies^2^ and Stress Granules^3^ as well the nuclear Cajal Bodies^4^ and Nuclear Speckles are some paradigmatic example of functional MLOs in the cell. RNA and intrinsically disordered proteins (IDP) are the major constituents of biomolecular condensates in physiological conditions.^5^ The inherent disorderedness of IDPs coupled with their low sequence complexity facilitates multivalent inter-molecular interactions leading to formation of these droplets (condensates).^6,7^ The most commonly studied phase separating IDPs with low complexity domains (LCD) include RNA binding proteins (RBPs) such as Fused in Sarcoma (FUS), heterogeneous nuclear ribonucleoproteins (hnRNPs) and TDP-43.^8–10^ These proteins are preferentially enriched with amino acids such as Tyrosine, Phenylalanine, Glycine, Serine, Arginine, Glutamine and Asparagine.

Several work have shown that the LCDs of these proteins form soluble and reversible aggregates composed of amyloid-like fibrils, ^11–14^ which are structurally and thermodynamically distinct from the insoluble and irreversible amyloid fibrils associated with protein misfolding of *α*-synuclein and A*β*.^15,16^ Essentially, the irreversible amyloid core consists of cross-*β* motifs in several layers forming tightly inter-digitated interactions. There is a body of evidence that now show that deviations in beta strands from the tight pleated conformation leads to functionally relevant fibrillar condensates that are reversible under physiological conditions.^12^ Certain mutations in LCDs could rob this reversibility as observed in neurodegenerative conditions such as Alzheimer’s,^19^ Parkinson’s,^20^ Amyotrophic lateral scleroris, Multisystem proteinopathy and amyloidosis. ^9,21^

In pursuit of understanding this functional assembly driven by IDPs, David Eisenberg and co-workers have recently identified short motifs within the LCDs of RNA-binding proteins that are capable of mediating reversible fibrillization. These sequence motifs are now identified as LARKS (low-complexity aromatic-rich kinked segments)^18^ and EAGLS (extended amyloid-like glycine-rich low-complexity segments).^17^ X-ray crystallography and diffraction studies of LCD containing proteins like hnRNPA1,^18^ FUS,^18^ TDP-43^22^ and NUP98^18^ revealed that when LARKS were isolated as peptides, they formed a cross-*β* structure containing kinks in the beta strands mostly at Glycine or aromatic amino acids.^18^ To identify potential LARKS in the human proteome, the same group developed a method of computational 3D profiling (LARKSdb) protocol that detects the regions in protein resembling LARKS by assessing the compatibility of query sequences with a template structure and assigning Rosetta energy score.^18,23^ Using this prediction tool, Eisenberg’s group has performed a genome-wide search of LARKS across different organisms and found these motifs to be abundant in Homo sapiens, Drosophila melanogaster, Mycobacterium tuberculosis, and Escherichia coli, which corroborate the notion of LARKS as an evolved structural motif.^24^ The assembly of LARKS-containing peptides form reversible fibrils that are transiently stabilized by weak interactions since the presence of kinks prevent the side chains from interdigitation across the *β*-sheet interface. This results in reduced buried surface area compared to the disease-causing amyloid fibrils. Extended *β*-strands, characterized by nearly linear conformations, is also associated with the reversible nature of amyloid formation. This ability of LARKS sequence motifs to deviate from classical pleated-*β* strands organization presumably contributes to the reversibility of the aggregation. The body of research highlights the importance of understanding the structural nuances of amyloid fibrils in the context of reversible self-assembly processes. ^17^

The amyloid aggregation process is complex, involving the formation of various metastable aggregates, ranging from small soluble species like dimers to nonfibrillar oligomers with some *β*-sheet structure.^25^ Our understanding of the nucleating factors and thermodynamic/kinetic landscape of fibril formation and the associated pathogenesis, both at structural and cellular levels, is continuosly evolving in recent years. ^26–28^ Particularly, early onset and the the role of oligomers in aggregation is debated, with conflicting evidence on whether they are on or off the pathway to amyloid fibril formation and their involvement in amyloid diseases. ^29^ Some studies suggest that pre-fibrillar aggregates are key contributors to cellular toxicity with the toxicity linked to their structural properties rather than their amino acid sequence. ^30^ In contrast, mature fibrils are considered largely non-toxic and may protect cells by sequestering harmful oligomers.^31^ Overall, the structure-to-function mapping of the biomolecular fibrillar assembly has been an area of active research and a better clarity is needed over the driving ingredients of reversible condensate formation at a structural level. One of the major deterrents in this exploration is unambiguous structural characterization of esoteric motifs such as beta-kink that are central to reversible fibril condensate formation. Parameters that accurately assign kinked-beta motif could be used as reliable reaction coordinate(s) to investigate the fibril formation pathways. The primary bottleneck in accurate characterization of beta-kink from given atomic coordinate data-set is due to the availability of limited high-resolution experimental structures. This is where Physics-based molecular dynamics (MD) simulations can contribute as a generative model to produce high quality conformational data sets. We use all-atom simulation data to reliably generate high-resolution conformational ensemble of conformations for IDPs that are known to have stretches that can form beta-strands. Along with the experimentally available data, these simulation-created structures can provide us with an exhaustive dataset to train algorithms that accurately characterizes motifs such as beta-kinks. This, in turn, gives us a better chance at investigating the features of condensates at single and multimer levels in a statistically rigorous manner.

In this work, we combined simulation-derived structures and experimentally available dataset to focus on the various structural features of *β*-strands in fibril-forming reversible and irreversible condensate systems. Among the different types of existing *β*-sheet motifs (pleated, kinked and extended sheets), we have focused on the non-trivial kinked-*β* sheets.

Aside from the existing framework to define the kinked-*β* sheets,^17,18^ we have recruited another set of parameters called bend angle (*θ_B_*) and rotation angle (*θ_R_*) that we find distinctly classifies kink and non-kink motifs. In 2015, Fujiwara and co-workers have prescribed *θ_B_*-*θ_R_* angles to accurately define the structural aspects of *β*-barrel proteins. Incidentally, use of this set of structural parameters led us to a faithful annotation of kinked-*β* sheets that we discuss in details below. We use support vector machine, a supervised machine learning technique, to specifically separate the obvious conformational kinks among the generated simulation dataset. We validate our classification map using the available solved structural dataset in protein data bank. Using our protocol on an extensive generative dataset from high-resolution atomic simulations on IDPs such as LCD of hnRNPA1, *α*-synuclein and a*β*-42, we also provide a library of potential functionally active kinked motifs that can be used for screening unresolved kinked datasets in new studies.

The remainder of this article is structured as follows. We elaborate our key findings in the result section. Following the Results section, we have a brief "Discussion" section to summarize the scope and future utility of our work. In the "Methods" section, we describe our protocols in details and discuss how we have used molecular simulations as a generative tool to create conformation dataset that helps us in developing the secondary structure assignment algorithm for kinked-*β* sheets. We also provide definitions of the structural parameters *θ_B_* and *θ_R_* used for training and explain the attributes of supervised machine learning based classification. Additionally we present details about the downstream analyses used to compare the efficacy of the various structural parameters mentioned in this paper. All our data, including simulation trajectories, the model used for supervised classification, and the associated codes, are available online for public access.

## Results

### *θ_B_* − *θ_R_* feature space unambiguously assigns kinked beta motifs

In order to structurally capture and quantify the kinked-*β* strand segments, our first objective was to establish one or more collective variables that can optimally define the motif using the coordinates data. For this purpose we chose *θ_B_* (bend angle) and *θ_R_* (rotation angle) as our parameters Fig. 1 (a,b).^32^ We have provided the elaborate formalization of these parameters in the Methods section. Briefly, in the Fig. 1 (a,b), the white circles denote four consecutive *C_α_* atoms from four consecutive residues. L,M and N respectively are the midpoint of the vector joining *C_α_*(*i*)-*C_α_*(*i*+1),*C_α_*(*i*+1)-*C_α_*(*i*+2) and *C_α_*(*i*+2)-*C_α_*(*i*+3). *θ_B_*, the bend angle, is defined by the angle between the two vectors **LM** and **MN**. **R***C_α_*(*i*+1)is a perpendicular from the point *C_α_*(*i*) on **LM** vector. On the plane (denoted by dashed ellipse) lying perpendicular to **LM** vector, vector **u** is a projector vector of **MN**. *θ_R_*, the rotation angle, is defined by the angle between the vector **R***C_α_*(*i* + 1) and the projection vector **u** (Fig. 1 (b). In Fig. 1 (b), the dashed circle below is a 2D representation of the rotation of vector **u** around the **R***C_α_*(*i*+1) vector where vector **LM** is the rotation axis.^32^ We apply this feature space on our training data set, which is sourced from more than 100 microseconds all-atom simulation data of four different proteins (hnRNPA1-PrLD, hnRNPA1-RGG, *α*-synuclein, a*β*42).^33–36^ We elaborate the process in the Methods section as well as in the Supporting Information (SI). In essence, we take a four-residue sliding window for every protein conformation in our training data, calculate *θ_B_* and *θ_R_* for the stretch and project the values on the *θ_B_*-*θ_R_* map. When projected on the *θ_B_*-*θ_R_* map (1(c)), we observe two considerably distinct clusters that occupies distinct regions comprising of kink and non-kink motifs. We used this projection to supervise our support vector machine SVM model for the final classification (Fig.1(d)). The SVM algorithm on our training data set generates a dividing hyperplane that faithfully classifies beta sheet motifs as kinked or non-kinked in the *θ_B_*-*θ_R_* map. In Fig. 1(d), the Yellow region is classified region as kink and the Blue region is non-kink. From the SVM statistical partitioning, the Green region is the region of ambiguity which is bordered by White lines that also provides us with the decision boundary. The color map depicts the posterior probability of the SVM. The advantage of this classification is clear when one compares the assignment using DSSP for a given conformation. For clarity, we show one such protein conformation in Fig. 1 (e) where the canonical *β*-strand is shown in Yellow and kinked beta-strand is shown in Red. Left panel of Fig. 1 (f) highlights the inability of the conventional secondary structure assignment tools such as DSSP^37^ to assign *β*-kinks. The Right panel shows the assignments using *θ_B_*-*θ_R_* values and the region marked in Red (residue stretch of (18 − 22)) is the kinked-*β* captured by our algorithm. Fig. 1 (g) provides the same information as 1 (f) but for a window of four-residues put together. The *θ_B_* and *θ_R_* angles are defined in a window of four residues. In other words, the nth window denotes the residue n, n+1, n+2 and n+3. For example, the 18*^th^* and 19*^th^* markers in Fig. 1 (g) are the corresponding markers of the two windows lying in 18 − 22 residue range in (f)). Likewise, 44 is the marker of the window lying in 44 − 47 window range. As for 66 – 67 residue range in (f)), (which consist of three residue) there is no window marker as a window consists of four residues.

**Figure 1:**
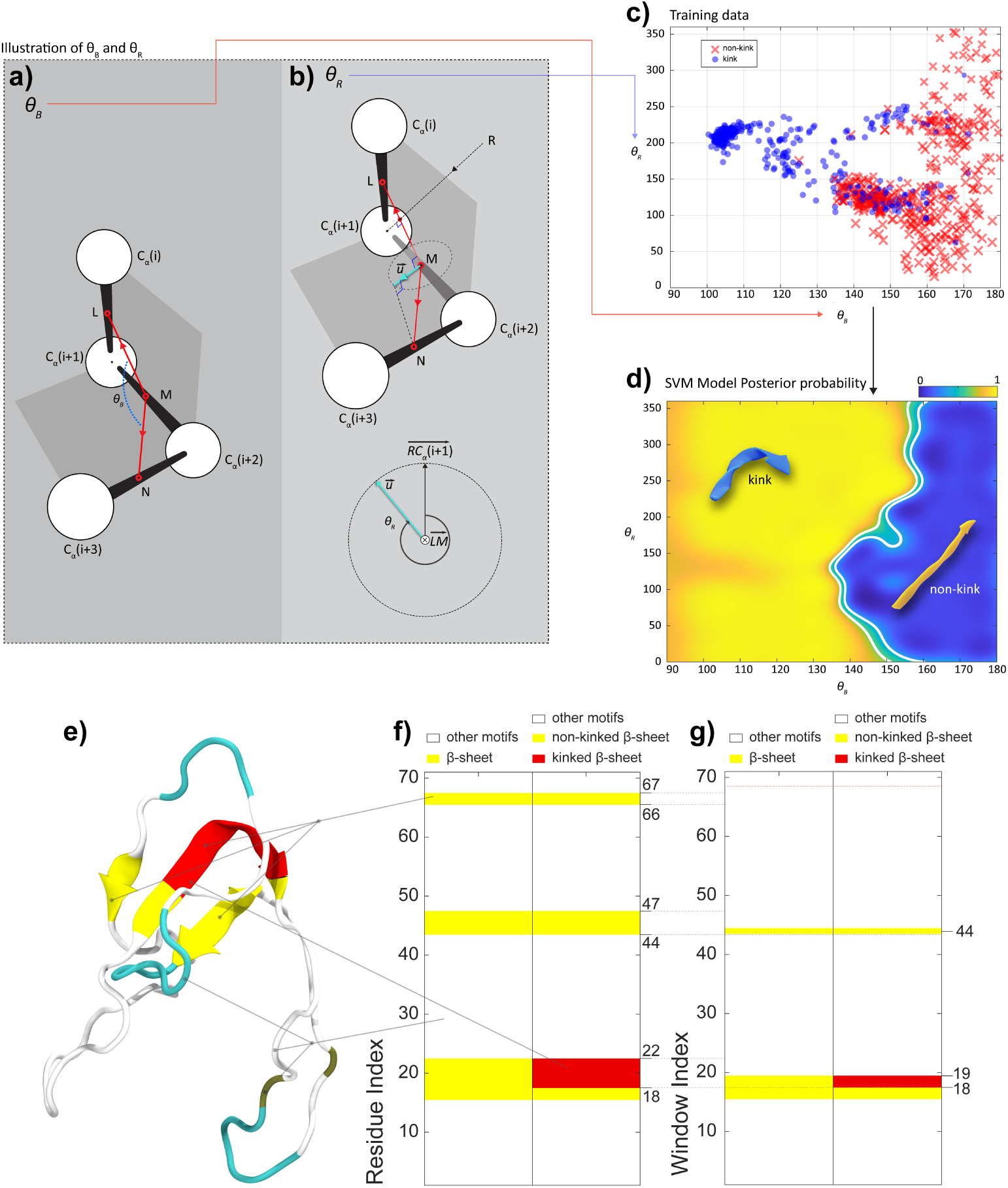
a,b) Schematic of *θ_B_* and *θ_R_* stereochemistry. c) Training data set used for the SVM classification represented in *θ_B_*, *θ_R_* space. Blue markers indicate kink and Red markers indicate non-kink beta sheets. d) The classification SVM model based *θ_B_* and *θ_R_* angles of training dataset using supervised learning with the dividing hyperplane. e) Snapshot of an example protein conformation used from the training data set. f) Comparison of DSSP and *θ_B_*-*θ_R_* algorithm for secondary structure assignment of kinked beta regions. g) 4-residue window-wise representation of 1(f), where each window index corresponds to the the residue with the same index and the indices of next three residue. In other words, the nth window denotes the residue n, n+1, n+2 andn+3. For example, the 18*_th_* and 19*_th_* markers are the corresponding markers of the two windows lying in 18 − 22 residue range in (f)). Likewise, 44 is the marker of the window lying in 44 − 47 window range. As for 66 − 67 residue range in (f)), (which consist of three residue) there is no window maker as a window consists of four residues.

In parallel, we wanted to examine the performance of other structural parameters with respect to the definition of kinked *β*-sheet motifs used in recent classification work for multiple types of *β*-sheets from Eisenberg group.^17^ Although there has been clear separation of extended *β*-sheets using some of these structural collective variables used in this work (*ψ*-*ϕ* angles, *C*1 − *C*4 distance, side chain torsion angle), none of these could clearly distinguish kinks from the set of *β*-sheet due to presence of notable overlaps among the kink and non-kink domains for a single parameter space. We have demonstrated these facts in details using our training data set as shown in Fig. S1-S3 in SI. We show that use of *θ_B_*-*θ_R_* has brought out a separate branch of kinked data set that is absent while using *C*1 − *C*4 distance-*θ_B_* or *C*1 − *C*4 distance-*θ_R_* collective variable pairs. We would like to note that our chosen parameters, *θ_B_* and *θ_R_* when implemented individually, does not perform any better than the mentioned structural parameters in ameliorating the ambiguity. However, the usage of pairwise structural parameters fair much better in terms of classification. In Fig.2 a), b) and c), we have shown the distribution of pre-labelled training dataset in the context of structural parameters *θ_B_* vs. *θ_R_*, *θ_B_* vs. *C*1 − *C*4 and *θ_R_* vs.*C*1 − *C*4 space, respectively. We have not used the *C* − *N* − *CA* − *CB* torsion angle as it doesn’t provide information about kinks involving Glycine leading to loss of information. Please see Fig. S2 in SI for details. In the insets of Fig. 2 a), b) and c), we show domain wise labelling of the 2D parameter space, where Blue is the domain occupied solely by kinks, Teal is the domain occupied solely by non-kinks and Yellow is the domain occupied by both i.e the region of overlap. To decrease the ambiguity, one should aim for decreasing the region of overlap. To quantify the performances of these combination parameters, we have computed three different metrics: Areal percentage overlap score, Domain percentage overlap score and min-Domain percentage overlap score. These scores are presented in Fig.2 d). The definition of these metrics can be found in Methods sections. A comparison of these metrics for three pairwise structural metrics show that the combination of *θ_B_*-*θ_R_* is performing better than the other two sets of parameters. As mentioned earlier, when visualised in a 2D space of *θ_B_*-*θ_R_*, these two parameters have been able to achieve a better resolution of the previous ambiguous regions into new branching subdomains which have reduced the ambiguity (which is not true for any other combinations of above mentioned collective variable). Please see the movie file (movie-1.mp4) in SI where the training data is projected on three different collective variables (*C*1 − *C*4 distance, *θ_B_* and *θ_R_*) and rotated around to show how the projections look when observed on two of the axes.

**Figure 2:**
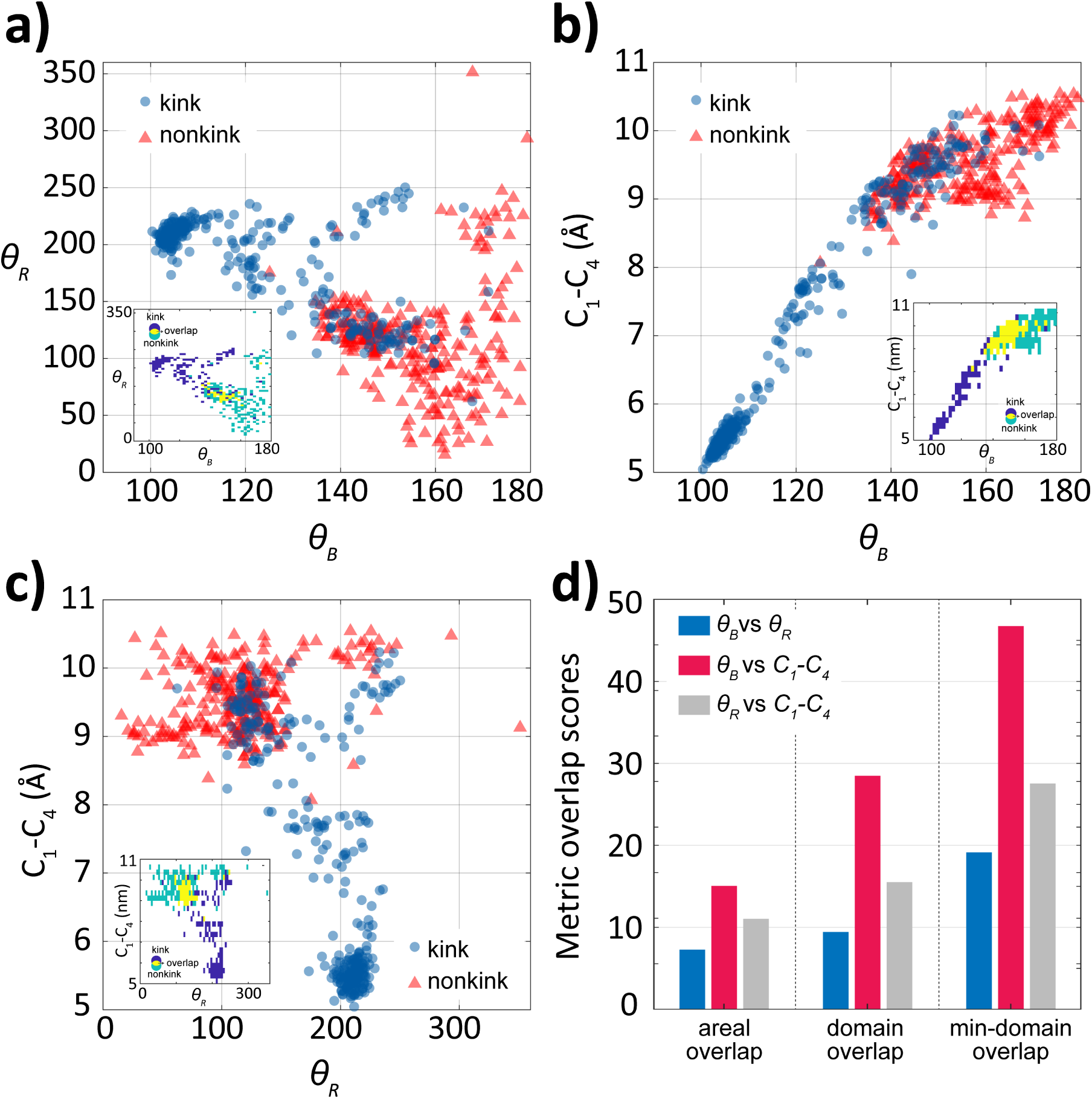
a), b), c) are the scatter plots of pre-labelled kink and non-kink training data set. Blue points are kinked beta-strand and red points are non-kinked beta-strand. a),b) and c) insets are the 2D histogram representation, where blue is the domain occupied by kinked beta-strands, yellow is the domain occupied by overlaps, teal is the domain occupied by non-kinked beta-strands. d) Bar plot showing three overlap metrics: Areal percentage overlap score, Domain percentage overlap score, min-Domain percentage overlap score respectively. Blue stands for the *θ_B_*-*θ_R_* pair, Pink stands for the *θ_B_*-*C*1 − *C*4 pair and Grey stands for the *θ_R_*-*C*1 − *C*4 pair.

### Experimentally resolved ***β***-kink structures honor ***θ_B_***-***θ_R_*** map

Our next objective was to corroborate our quantification technique using experimentally available fibril structures.To observe the performance of our trained SVM classification model, we took the total of 26 reversible and 119 irreversible experimental solved fibril dataset and calculated their *θ_B_* − *θ_R_* angles. We have made lists of irreversible and reversible fibrils and provide them as a MS Excel file in SI (ListSolvedFibrils.xls). Upon implementation of the classification model on these angles, we could segregate the fibrils into kinked and non-kinked beta stranded groups where Fig.3 a) is for the reversible fibrils and Fig.3 b) is for the irreversible fibrils. This validation was a significant show of the usability of the classification model.

**Figure 3:**
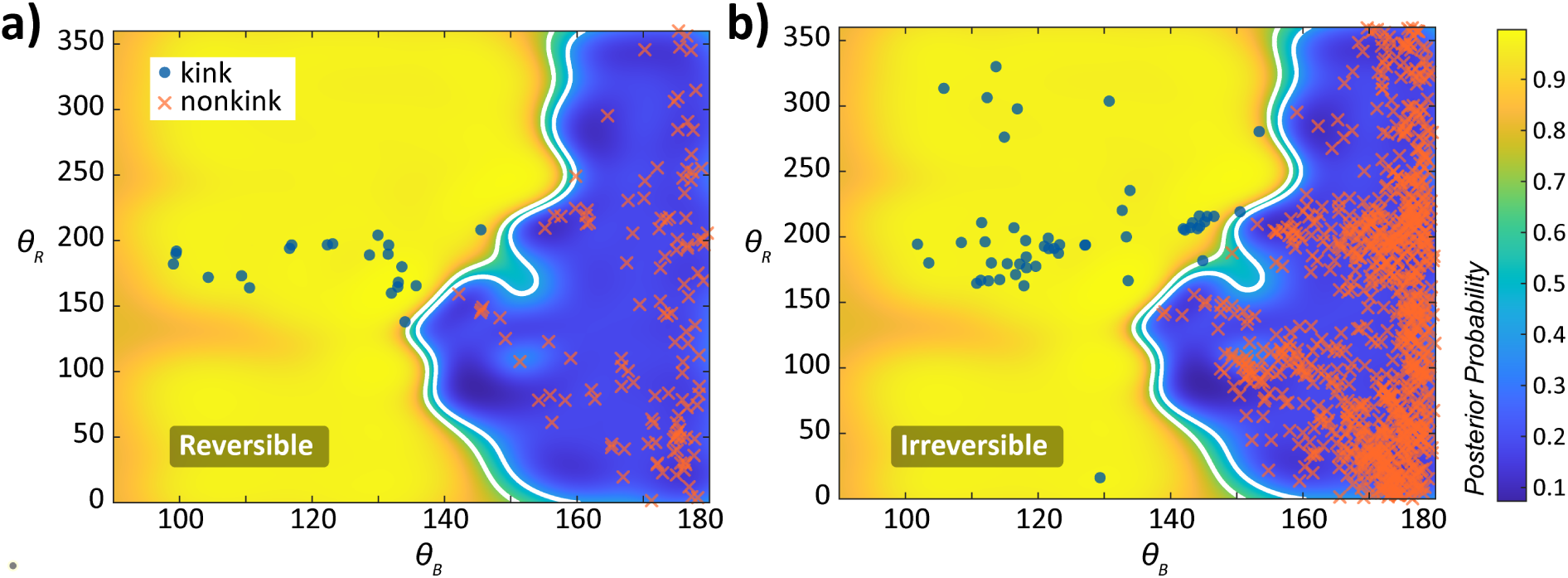
*θ_B_* − *θ_R_* based SVM classification map of a) reversible fibril dataset and b) irreversible fibril dataset. The yellow region is classified region as kink, the blue region is non-kink. The green region is the region of ambiguity which is bordered by white lines. This also creates the decision boundary. The color map depicts the posterior probability of the SVM.

### *β*-kinks appear frequently in simulation-generated IDPs conformations

Bolstered with this evidence in hand, we went ahead to implement the model on our simulation dataset. The execution of SVM on RGG-box (50 amino-acids) and PrLD region (70 amino-acids) of hnRNPA1-LCD simulation data provided us with classified *θ_B_* and *θ_R_* angles dataset across all the *β*-strands in terms of kinked and without-kink *β*-strand motifs Fig.4. The folders containing the trajectory files are available on the following Figshare repository: https://figshare.com/s/ea0cfee10359936eb3e0. The corresponding movie files of the trajectories for RGG-box and PLD domain of the hnRNPA1-LCD are available as MP4 files, movie-2.mp4 and movie-3.mp4, respectively in the the same Fighare link. In the Fig. 4a) and b), we could see the separation of *θ_B_*-*θ_R_* space in Yellow (kink) and Blue(non-kink) classes where the white line has been acting as decision boundary. Along with this, we were able to locate the significant occurrence of kinks among all the existing *β*-strands across the entire PrLD and RGG-box sequence over a 1 microsecond time frame (Fig. 4c) and d)). In both 4c) and d), we have kept the time frames and residue windows on X and Y axis respectively. At different snapshot we have noticed various residue windows are getting flagged for containing a kink in the *β*-strand of the associated segment. We have also taken available all-atom simulation data of *α*-synuclein and amyloid-*β* from Robustelli and co-workers^35^ and Vendruscolo and co-workers^36^ and implemented our classification model on the respective trajectories (SI fig 4 a), b)). Here as well, we were able to see categorization of the existing beta-strands into kinked and non-kinked groups. Along with the abundance of kinks all over the course of the simulation trajectory, we could also observe the relative permanence of some of the motifs while the others had a more transient life span. This robustness of the kinks among certain regions of the protein may be translated into the functional significance of the residues promoting a more persistent kink.

**Figure 4:**
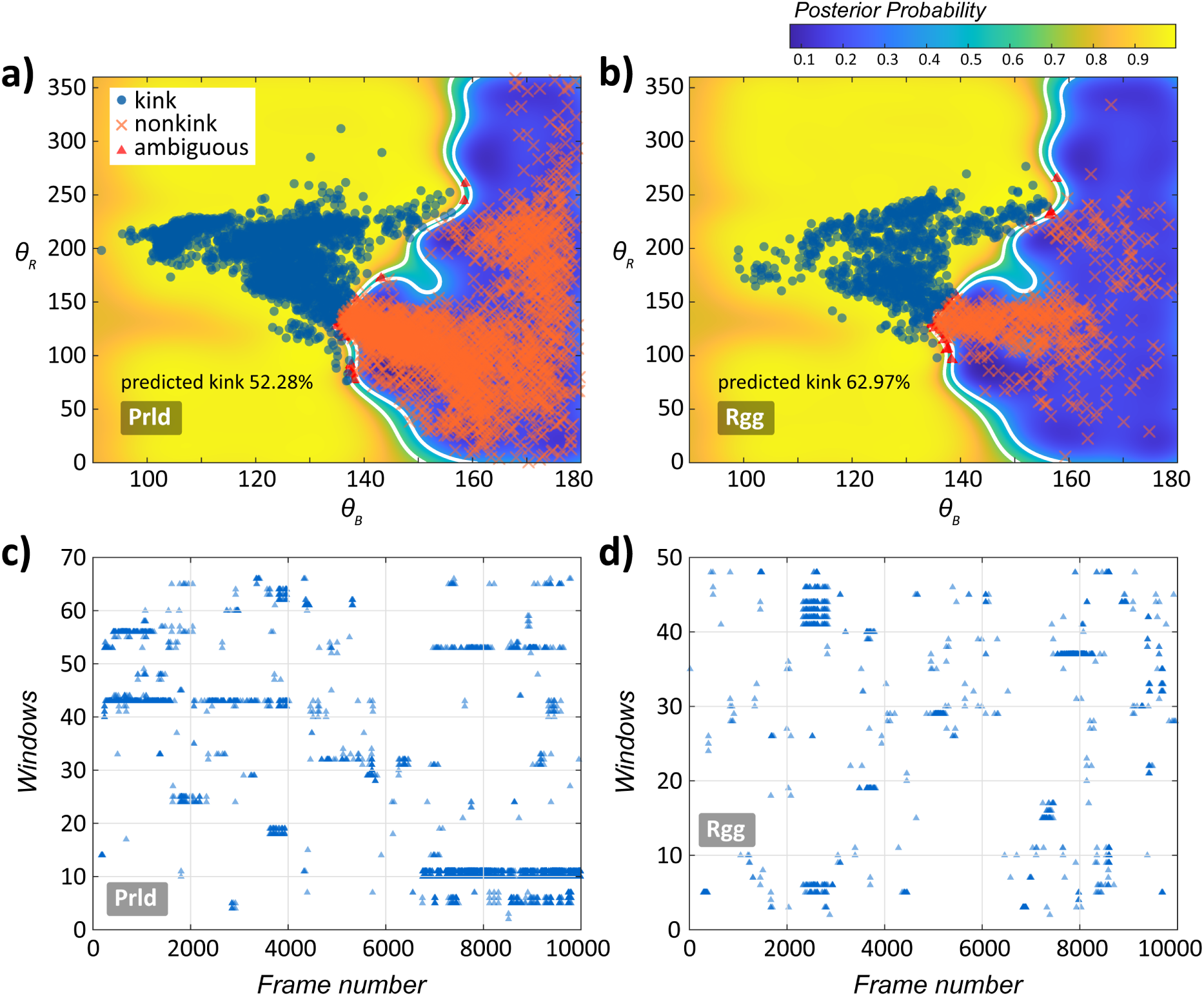
Kinked-*β* strands classified based on *θ_B_* and *θ_R_* angles using SVM for a) PrLD ensemble b) RGG ensemble. Blue (orange) markers on the left (right) side of the boundary stand for presence(absence) of kinked-*β* strands. The markers flag the four residue frames across time and sequence which form the kinks. The color map indicates the probability of kink or non-kink spanning from yellow to blue (kink to non-kink) with boundary at 0.5 (green). The white borders encompass the region of ambiguity. Kinked-*β* strands profile plot with respect to time evolution for c)PrLD and d) RGG. On X-axis and Y axis, time and residue windows are present, respectively. From a 1*second*trajectory, we have taken every 100*_t_h*frame. The markers flag the windows across time and sequence which form the kinks.

### Kinked-***β*** motifs scatter in and around the allowed regions of Ramachandran plot

Followed by this, we wanted to observe the distribution of kinks over the Ramachandran’s *ϕ* − *ψ* landscape. We started with visualising our training data set in *ϕ* − *ψ* space (see SI. Fig. S5). We see that a statistically dominant percentage of the non-kinks are occupying the known *β*-strand region of the Ramachandran map. However, the kinked motifs are spread in multiple regions of the map indicating a wider reach over the *ϕ* − *ψ* space. Despite this specific distribution of kinks in the *ϕ* − *ψ* space, the distinction based on the *ϕ* − *ψ* angle are not enough as we observe that a considerable number of the kinked points share space with the known *β*-strand region. This in turn renders difficulty to the classification only in terms of *ϕ* − *ψ* space. Here we again emphasize the implication of *θ_B_*-*θ_R_* in considerable reduction of the ambiguity between kink and non-kink motifs.

The inspection of *ϕ* − *ψ* angles of the kinked motifs in hnRNPA1-PrLD (Yellow triangle) and RGG (Red triangle) ensemble provided us with a similar Ramachandran space where we could again see similar spread on the *ϕ* − *ψ* space Fig.5(a) which are different from the strict *β*-sheet signature location. We have also highlighted those *ϕ* − *ψ* where Glycine is a contributing residue ( plus as PrLD Glycine and circle as RGG Glycine) to show the spread of this particular residue. Fig.5(a) clearly indicates the deviation of kinked *β*-motifs from standard *ϕ* − *ψ* regions while adhering to the general allowed region of the Ramachandran plot which in turn certifies the existence of this type of motif in protein conformations at large. Along with the simulation data set, we have also taken up the reversible (Fig.5(b))and irreversible fibrils (Fig.5(c)) experimental data set^17^ and showed that the *ϕ* − *ψ* angles of the kinks obtained in the experimental datasets belong among the already observed clusters in the Ramachandran space. Overall, in this part, we have shown the significance of *θ_B_* and *θ_R_* in capturing the kinked structures and their arrangement in the Ramachandran conformational space.

**Figure 5:**
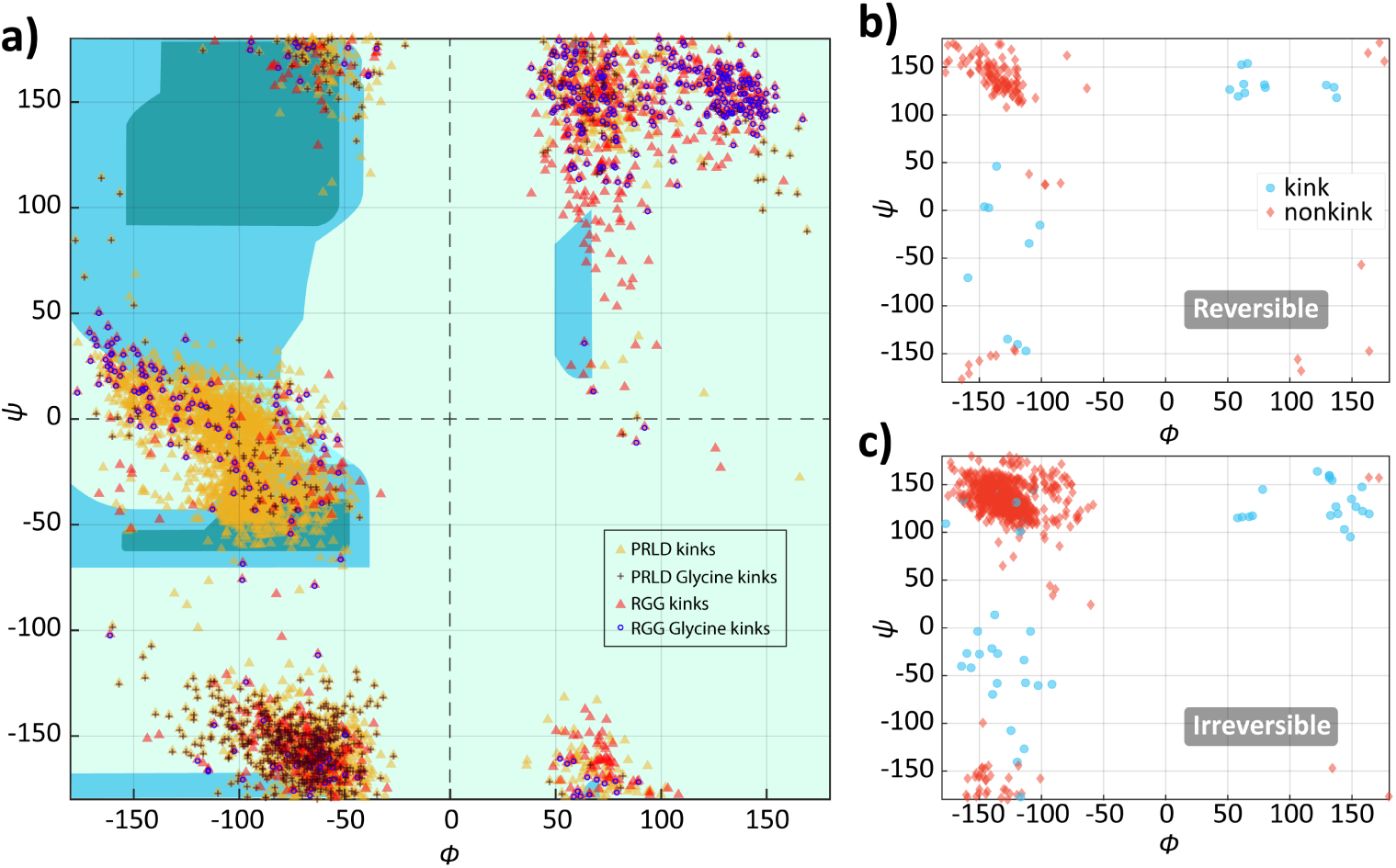
a) Distribution of kinked beta motif points in the Ramachandran phase space. Yellow triangles denote all the kinked beta-strand motifs from PrLD simulation. Black plus indicates the subset of kinked beta motifs from PrLD which include Glycine in the four residue window. Red triangles denote all the kinked beta-strand motifs from RGG simulation. Circles indicates the subset of kinked beta motifs from RGG which include Glycine in the four residue window. The teal and indigo colored regions are the areas designated to standard secondary structures of a protein. b)The *ψ*-*ϕ* angles distribution of kink(blue) and non-kink(red) data from irreversible fibril dataset. The list of experimentally solved irreversible fibrils are collected from the work done by Prof. Eisenberg’s group^17^ c) The *ψ*-*ϕ* angles distribution of kink(blue) and non-kink(red) data from reversible fibril dataset. The list of experimentally solved reversible fibrils are collected from the work done by Prof. Eisenberg’s group.^17^

### A subset of kinked-*β* strands form LARKS

Along with the structural features, the LARKS captured by Prof. Eisenberg’s group^12^ have specific sequence signatures. In order to integrate these residue compositional information into our kinked-*β* strand definition and thereby extract the LARKS subset, we developed a sequence filter for each four-residue window given in the Table 1. Applying these sequence filter in a ‘residue-wise’ manner on the kinked-*β* strand dataset we could distill a section of kinked motifs which can be characterised as LARKS. We have shown this LARKS subset in the *θ_B_* and *θ_R_* phase space of SVM classification map where green triangles are PRLD simulation data and pink circles are from RGG ensemble (Fig. 6 a)). In SI Fig. 6 a) and b), the LARKS content of residues from the PrLD and the RGG ensemble has been plotted for 1*µS* dataset. A comparison within the LARKSdb predicted data set and our calculation (Fig. 6 (b) (c)) shows more number of residues assigned as LARKS-forming residues by our method. Followed by checking for LARKS in the simulated ensembles, we have also checked for kinked beta-strands in experimentally solved LARKS fibril crystals obtained from Prof. Eisenberg’s group.^12^ Upon implementing our classification model on the coordinates of six LARKS fibrils, we could capture the kinks in all 6 systems with varying numbers (Fig. 6(d), SI fig. 7 a)) and showed those points in the SVM map of *θ_B_* and *θ_R_*. Additionally, we wanted to see the result of implementing the sequence filter on this kinked-angle dataset. Using the sequence signature mentioned in Table-1 we have captured the specific LARKS segments in these experimental dataset (SI fig. 7 b)). The peach color tiles show presence of *β*-kinks in a sequence verified peptide fragment. Important point to note is, in Fig.6 (c), we could successfully capture one of the experimentally proven LARKS (red highlight) existing in hnRNPA1-^243^*GY NGFG*^248^ (PDB ID: 6BXX) peptide segments. As a whole, our method is able to recognise the irregular *β*-strand structure in a tangible manner for all six experimentally solved LARKS fibrils. LARKS has been shown as functional units to induce reversibility in amyloid fibrils. Overall in this study, we are able to show that among the broader set of kinked-*β* structure, LARKS is a smaller section of sequence specific motifs containing the similar irregular *β*-motifs.

**Figure 6:**
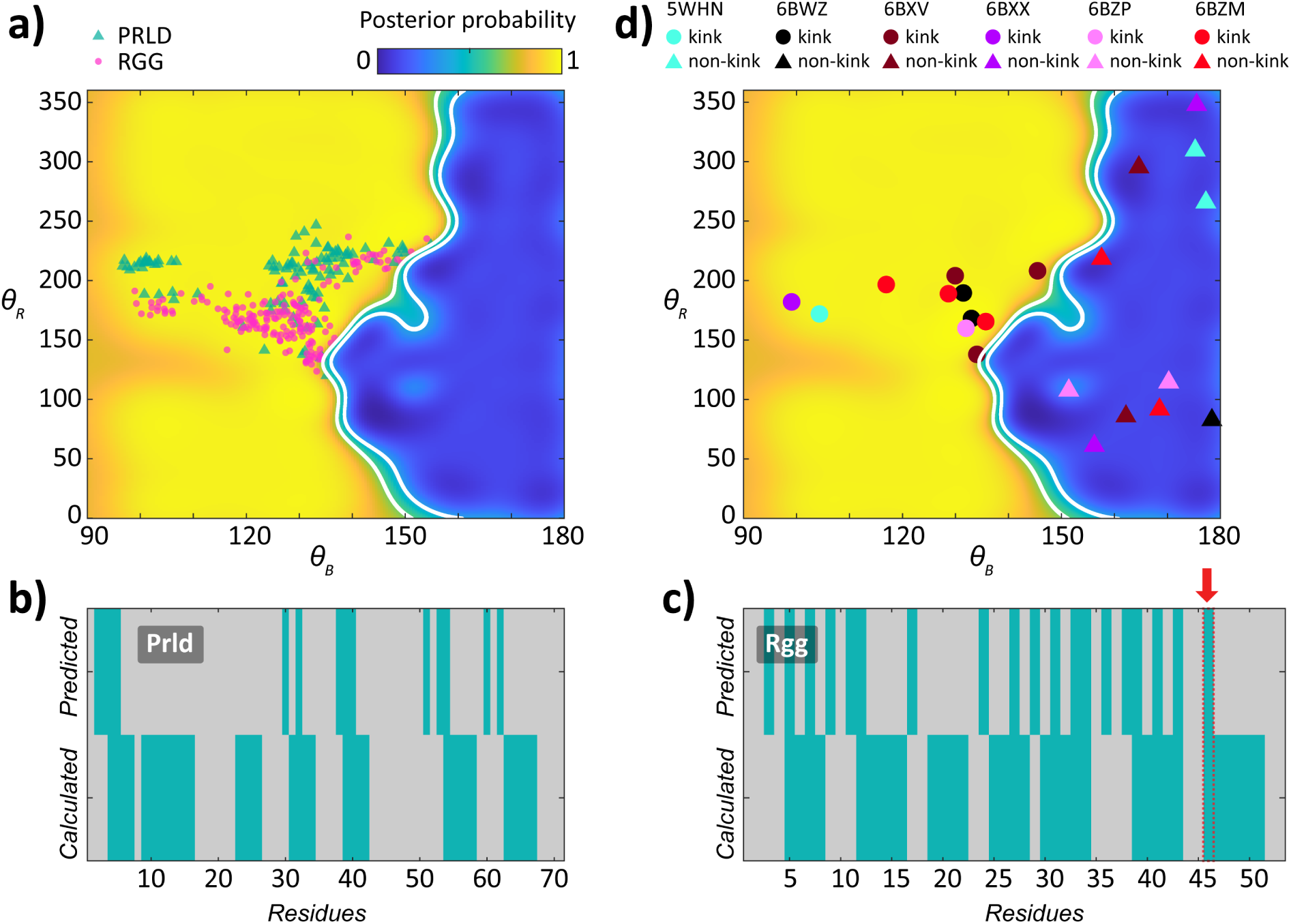
a) *θ_B_* and *θ_R_* angles phase space corresponding to LARKS motif for PrLD and RGG ensemble. Green triangle are from PrLD ensemble, Pink circles are from RGG ensembles. The color map indicates the probability of kink or non-kink spanning from yellow to blue (kink to non-kink) with boundary at 0.5 (green). b) and c) Comparison between LARKS prediction by LARKSdb and LARKS calculation by our method for PrLD and RGG ensemble respectively. The top panel is for predicted dataset. The bottom panel is for calculated data set. The cyan tiles stand for residues participating in LARKS formation. The grey tiles stand for residues having no contribution in LARKS formation. The red highlighted region in (c) has been found in experimentally solved fibril as well. d) *θ* − *B* and *θ* − *R* angles profile of kinked-*β* strands in experimental data. This plot shows the position of *θ* − *B* and *θ* − *R* angles of the kinked-*β* strands in the classification domain obtained from 6 experimentally solved crystal structures showing LARKS: TDP43-^312^*NFGAFS*^317^(5WHN, cyan) showing kink but sequence doesn’t match LARKS definition, FUS-^37^*SY SGY S*^42^(6BWZ, black), FUS-^54^*SY SSY GQS*^61^(6BXV, maroon), hnRNPA1-^243^*GY NGFG*^248^(6BXX, violet), FUS-^77^*STGGY G*^82^(6BZP, pink), nup98-^116^*GFGNFGTS*^123^(6BZM,red).

**Figure 7:**
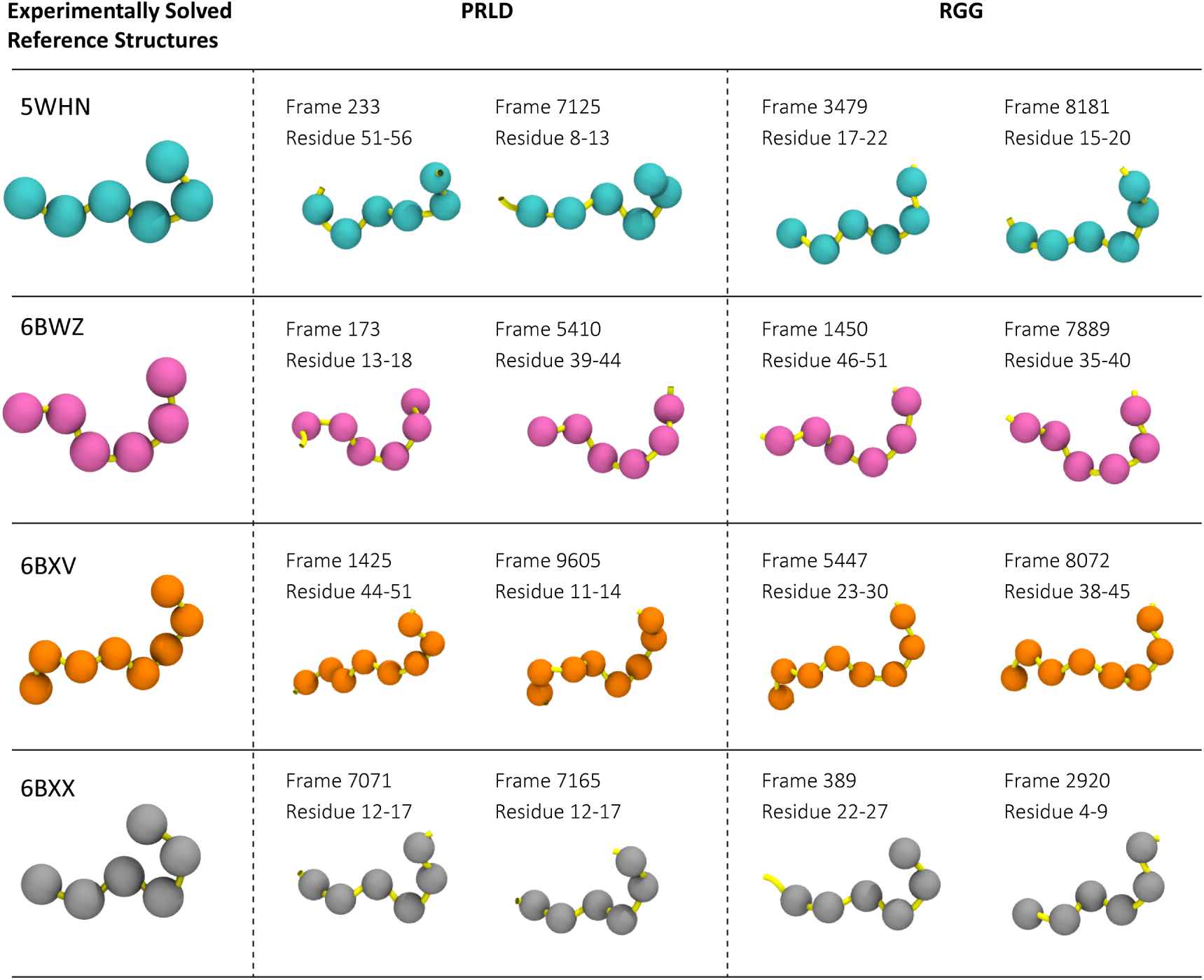
Column 1 shows the *C_α_*-atom representations of the experimental fibril data used for building the library (TDP43-^312^*NFGAFS*^317^(5WHN), FUS-^37^*SY SGY S*^42^(6BWZ), FUS-^54^*SY SSY GQS*^61^(6BXV), hnRNPA1-^243^*GY NGFG*^248^(6BXX)). Column 2 shows two of representative kinked beta-strand segments obtained from the PrLD ensemble which match the corresponding experimental data. Column 3 shows two of representative kinked beta-strand segments obtained from the RGG ensemble which match the corresponding experimental data. The frame number and residue range for the segments corresponding to respective trajectories are mentioned in the table. The RMSD threshold used for this study is 0-1.5Å.

**Table 1:** A four residue window sequence filter for extracting LARKS.

| Residue 1 | Residue 2 | Residue 3 | Residue 4 |
| --- | --- | --- | --- |
| [SNQGYF] | [GS] | [YF] | [SNQGYF] |
|  | [YF] | [GS] |  |

### Predicted beta-kink peptide library for future screening

Finally, we have focused on the generative aspect of this SVM classification model. Despite the extensive nature of experimental work, the kinked beta-strand dataset is not statistically rich. To improve the available data bank, we used our simulation derived classification filtered data to bring forth a library of representative kinked beta-strand segments. In order to look for the prototypical structural motifs, we looked for the peptide segments from our simulation data (PrLD and RGG) which provided a close structural match with the experimental data. To this end, we used root mean square deviation (RMSD) as our evaluation criterion among experimentally solved crystal structures (reported comparison against four experimental systems:TDP43-^312^*NFGAFS*^317^(5WHN), FUS-^37^*SY SGY S*^42^(6BWZ), FUS-^54^*SY SSY GQS*^61^(6BXV), hnRNPA1-^243^*GY NGFG*^248^(6BXX)) and simulation dataset. A RMSD comparison taking 1.5Å as a systematic cutoff provided us with possible functionally significant sections among the entire kinked dataset for each experimental data (SI Fig. 8 and 9). In the SI Fig. 8 and 9 the circle markers (the marker colors 0-1.5Å refers to corresponding RMSD value) within the scatter plot indicates the experimentally similar peptide segments. Fig. 7 shows a table where the first column contains the experimental references used for these analysis, second and third column shows two example representative motifs for PrLD and RGG ensemble, respectively which match the corresponding experimental structure within a RMSD threshold of 0-1.5Å. We have provided the full library in the form of frame number and residue range using which all the library members can be extracted from the PrLD and RGG trajectory. Overall, through these set of analysis we have managed to provide a vast set of kinked motifs which act as representatives of kinked *β*-strand structures found across different proteins. This library can be used as a starting point for screening of more involved downstream experimental analysis in pursuit of finding functional significance.

**Figure 8:**
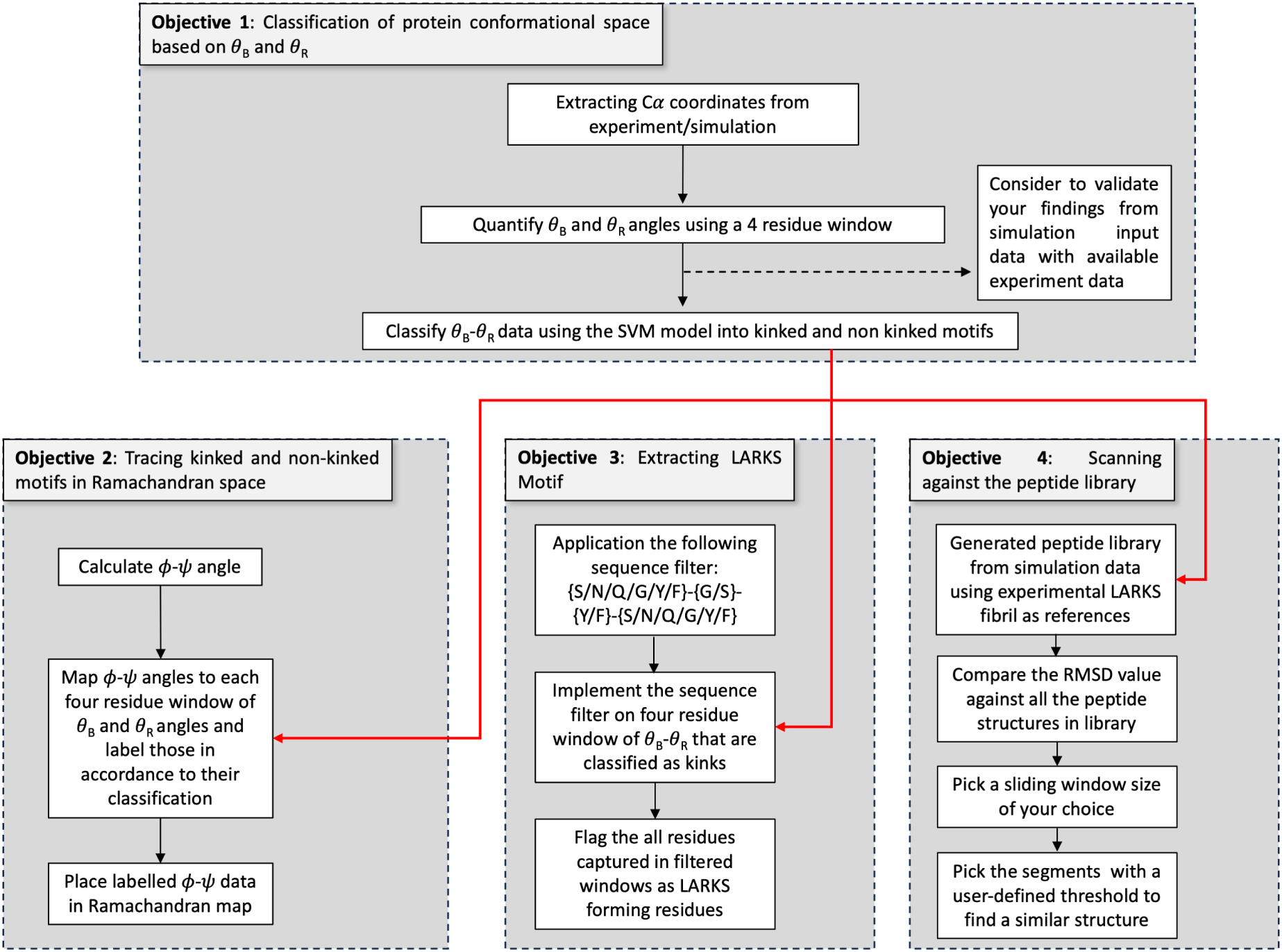
Flowchart describing the steps of the prescription of SVM classification (left panel), LARKS extraction (middle panel) and building and use of peptide library (right panel) in the paper.

**Figure 9:**
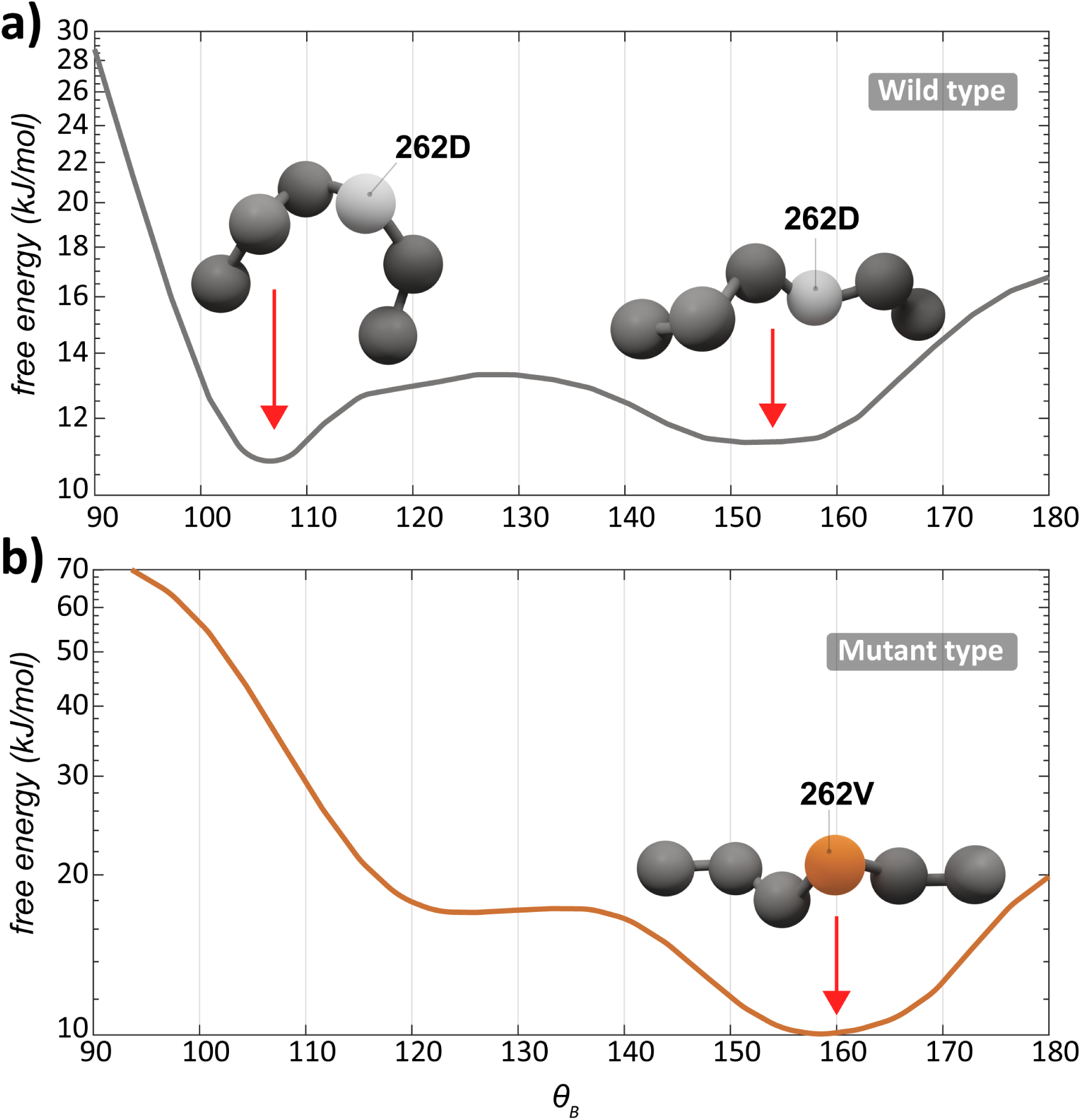
Free energy vs. *θ_B_* plot for a) wild type ^259^*SY NDFG*^264^ peptide segment and b) mutant ^259^*SY NV FG*^264^ peptide segment. The representative structures corresponding to the energy minimas are depicted as *C_α_* atom format in the plots. The mutant *θ_B_* population has reached a deeper minima compared to that of wild type. This is an indication that once the population reaches the minima, it favours the lower free energy well. The orange atom is the position of mutation.

## Discussion

Collectively, in this work, we provide a prescription of labeling kinks given a set of atomic co-ordinates. The flowchart of the entire work is provided in Fig. 8 where each block corresponds to the workflow and highlights findings. The strong suit of this classification prescription lies in its ability to capture the kinked beta motifs with significant accuracy than any other structural analysis previously reported. It should be noted that there is some ambiguity in the non-kink phase space where potential kink can be found sparsely among various non-kinked segments. As evident from the training data, there are two branches along which kink structure gradually evolves to non-kink with no hard boundary, and hence can not be resolved by the use of *θ_B_* and *θ_R_* or any other structural parameters already tried in the literature. Further research on possible parameters specific to kinked segments is required to alleviate this ambiguity. Nevertheless, among the existing measures, our technique has been able to eliminate significant amount of ambiguity (by resolving the two branches) in terms of distinguishing between kink and non-kink segments. We have also checked for the *β*-turn features in one of our test data set (hnRNPA1-PrLD). *β*-turn are identified using hydrogen-bond length and angle criteria stated by Venkatachalam et.al. (^38^). Interestingly, we find that all the residue windows satisfying the *β*-turn criteria fall within the kink dataset as shown in fig. SI10. As illustrated in this figure, the *β*-turn (Pink marker) occupies a small sub-domain of the kink-domain. *β*-turns are previously reported to take part in reversible fibril formation. Thus, *β*-turn being a small subset of all kink motifs further supports the idea of reversible fibril formation by kink motifs overall. Note that our technique expands on previously known beta motif (i.e. beta turns) that is reported in reversible fibrils by classifying the domain space via other structural classification and exploring the full domain space where reversible fibril formation is possible.

Additionally, as an exploratory attempt, we have also ventured into the future scope of the structural parameters *θ_B_* and *θ_R_*. Potential of mean force calculations are usually carried out as a function of collective variable (reaction cordinate) to extract functionally relevant conformations. Here we use *θ_B_* to inspect an established mutation D262V in the PrLD region of hnRNPA1 that is know to lead to fibril formation.^9^ Upon substitution of Aspartic acid by Valine in the steric zipper motif (^259^*SY NDFG*^264^), the reversible fibril changes it nature and tends to form solid irreversible fibrils leading to various neurodegenerative disorders.^9^

We have performed metadynamics simulations for both the wild type and mutant type of ^259^*SY NDFG*^264^ single chain segment using *θ_B_* as collective variable. We provide the implementation details of Metadynamics in the Methods section. We observed that wild type shows two free energy basins with a shallow barrier in between as an indicative of reversibility between kinked and pleated nature of *β*-strand (Fig. 9(a)). On the other hand, the mutant shows one major deep free energy minima near the *θ_B_* phase space of pleated *β*-strand pointing at increased stability of pleated *β*-strand and higher energy cost of conversion to kinked *β*-strand (Fig. 9(b)). This in turn is symptomatic of the irreversible fibril formation upon mutation. We show the convergence of our metadynamics calculations in the SI Fig. S11. In a nutshell, we have shown the future scope of these collective variables as a tool to investigate the nature and mechanism of gel-to-sol fibril switching.

In summary, creation of reversible membrane-less organelles are cell’s way of efficient functional compartmentalizing over space and time. Cells across different species have become adept in making such organelles given it’s economic nature of resource usage, especially during stressful conditions. Given the cross-species utility of this phase separation induced condensates, we decided to implement our kinked-*β* strands quantification method on the available experimental and simulation derived fibrils of different proteins found in various organisms. In this work we have shown the classified kink and non-kink information of experimentally solved segment of amyloid-*β* protein and of simulation obtained ensemble data. The existence of such kinked motifs in proteome of a diverse set of organisms indicates there significant functional importance spread over a broad genre of cellular activity including stress granule formation, creation of complex required for splicing, nuclear pore complex formation.

## Methods

### Simulation

We started with an initial unfolded structure of PrLD obtained from Iterative Threading ASSEmbly Refinement (I-TASSER)^39–41^ and to obtain the ensemble we have employed an advanced sampling technique called replica exchange with hybrid tempering (REHT) developed in our lab^33^ where both the temperature as well as the Hamiltonian across the replicas are changed. The force field used is Charmm36m.^42^ Table 2 contains detailed information of the PrLD system-setup. The system is solvated in a cubic box with a minimum distance of 1 nm from the surface of the protein using a 3-site rigid TIP3P water model. The system was also neutralized to maintain a physiological concentration of NaCl (0.15*M*). We used Gromacs-2016.5 patched with Plumed-2.4.1^43–45^ to carry out the simulation. We performed energy minimization of the solvated protein system using steepest descent algorithm for 50,000 steps to avoid any poor contacts. The energy minimized structure was then equilibrated sequentially in NVT and NPT ensembles for 1 ns.

**Table 2:** Advanced sampling system information of simulation for PrLD and RGG-box domain.

| Systems studied | Number of atoms | Number of replicas | Overall effective temperature range | Cumulative time per replica (in ns) | Average exchange acceptance probability |
| --- | --- | --- | --- | --- | --- |
| **PrLD** | 23538 | 24 | 300 K-500 K | 1000 | 0.3 |
| **RGG-box** | 22948 | 20 | 300 K-500 K | 1000 | 0.26 |

Next, the protein and the solvent were coupled separately to the target temperatures using the Nose-Hoover thermostat and the final production simulation was performed in the NVT ensemble. A cut-off of 1 nm was used for calculating the electrostatic and vdw interactions and Particle Mesh Ewald was used for long-range electrostatics. To integrate the equations of motions, the Leap-frog integrator with the time step of 2 fs was used. LINCS algorithm was used to constrain all the hydrogen atoms. Exchanges were attempted at every 1 ps interval.

In this approach, the Hamiltonian of the solute particle in the different replica is scaled-down up to 0.6 − 0.7 while also raising the temperature of the systems simultaneously up to a maximum of 500 K. This way, as a function of both lambda scaling and explicit thermostat conditions, a very high effective temperature is ensured to be realized on the protein solute to sufficiently overcome the energy barriers. The solvent is heated up mildly in order for the rapid reorientation of the hydration shell upon conformational change of the protein. The number of replicas used for all the systems are indicated in Table 2 and the corresponding temperature ranges and average acceptance probabilities are also indicated. Post processing analyses of all the trajectories were performed with the Gromacs analysis tools and the trajectories were visualized using VMD.

### Definition of ***θ_B_***-***θ_R_*** calculation

We defined a four residue window of a peptide with four consecutive *C_α_* atoms (marked as white circles Fig. 1 (a)^32^). L, M and N are the midpoints between *C_α_*(*i*)-*C_α_*(*i* + 1), *C_α_*(*i* + 1)-*C_α_*(*i* + 2) and *C_α_*(*i* + 2)-*C_α_*(*i* + 3) respectively. From *C_α_*(*i* + 1) point a perpendicular is drawn on the vector **LM** which we are calling as the vector *RC_α_*(*i* + 1). We designate *θ_B_* or bend angle of this peptide frame as the angle between the two vectors **LM** and **MN**. In order to quantify the rotation of *C_α_* atoms along the backbone of the peptide window, we define a rotation angle called *θ_R_* which ranges such that 0 *< θ_R_ <* 360. This is the angle between the perpendicular vector *RC_α_*(*i* + 1) and the projected component **u** of vector **MN** along a plane normal to vector **LM** (The plane is depicted as a dashed ellipse in Fig. 1 (b)). For visual clarity, in the Fig. 1 (b) bottom panel, we have also depicted the two vectors *RC_α_*(*i* + 1) and **u** (with the emphasis on the rotation angle *θ_R_*) along the above mentioned plane with **LM** vector pointing inwards to the page (Fig. 1(b)).^32^

### Classification of ***θ_B_***-***θ_R_*** using Support vector machine(SVM)

To classify the *θ_B_* − *θ_R_* angles, we have recruited a supervised machine learning based SVM classifier. We divided our simulation data set into three parts: training, cross-validation and test datasets. Training data consists of a list of *θ_B_* and *θ_R_* angle values for a window of four residue and their corresponding kink classification labels. This data set has been predominantly taken from the simulation PrLD dataset Fig.1 (c), and partially from RGG, *α* synuclein and a*β*-amyloid proteins. Cross-validation data is a combination of both PrLD and RGG ensemble data. The binary classification of kink and non-kink conformation of training is done based on visualisation of beta sheet motifs using STRIDE algorithm of VMD. For SVM classification we have used a Gaussian kernel function with Bayesian optimiser where the scores are weighted based on posterior probability values. The posterior probability(p) is shown as a colour map in Fig.1 (c) where *p >* 0.5 means the specific point falls within the kink class in *θ_B_*-*θ_R_* space with a corresponding probability p. Whereas, *p <* .5 indicates that the point falls in non-kink class with classification probability of 1 − *p* Fig.1 (c). This can be treated as confidence score for respective points. We have used the MATLAB package called fitcsvm^46^ for the classification purpose.

### Computing overlap metrics

In Fig.2 a), b), c) we have depicted the distribution of kink and non-kink motifs in different structural parameter-space namely *θ_B_* vs. *θ_R_*, *θ_B_* vs. *C*1 − *C*4 and *θ_R_* vs.*C*1 − *C*4. In all of these three 2D parameter-space kink and non-kink population share certain regions of overlap. The overlap is a combination **1)** of the span of the 2D domain where both types of motifs coexists and **2)** the population within such coexisting domains. For example, even if the size of a coexisting domain is significant, if the population within the domain is very less it would mean there is no significant overlap. On the other hand, even if a coexisting domain area is small but both kink and non-kink motifs densely populate the domain, the overlap in turn should be considered significant.

Here we introduce three metrics that quantifies the overlap in the 2D domains of *θ_B_* vs. *θ_R_*, *θ_B_* vs. *C*1 − *C*4 and *θ_R_* vs.*C*1 − *C*4. The areal percentage overlap metric considers both the population within the domain as well as the size of the domain. In areal percentage overlap score, we made separate 2D histograms of pre-labelled kink and non-kink training dataset into 2D bins for all three pairwise combinations: *θ_B_* vs. *θ_R_*, *θ_B_* vs. *C*1 − *C*4 and *θ_R_* vs.*C*1 − *C*4. Then for each combination metric, we took the minimum height between the kink and non-kink histograms in each bin which is basically the intersection between the kink and non-kink dataset at that particular bin. We also find the maximum height of the histograms within the said bin which gives us the union between two histograms at this particular bin. Followed by this we calculate the total union and total intersections by summing over all the bins. Next we implement Eq.1.

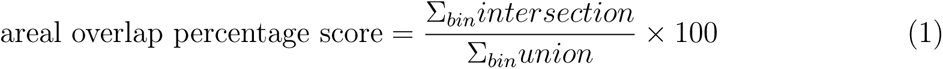

For the next two overlap metrics we only use relative size of the intersection domain to quantify overlap. In domain percentage overlap score, first we quantify the intersection domain area by the total number of histogram bins where both kink and no-kink motifs are present (i.e have a non-zero count for both kink and non-kink). We then calculate the union between the domains of kink and non-kink by counting the total number of bins where either motifs is present with non-zero count. This union is the quantification of the area of the domain that contains both motifs. Followed by this we are implementing Equation 2.

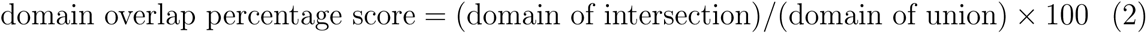

The min-Domain percentage overlap score is a variant of the above metric. Here, instead of taking the entire union-domain set as the denominator, we are taking the minimum contributing domain of the overlap as the denominator. The reason of choosing this specific metric is to tease out the true depiction of overlap in a relative manner. For e.g., in Fig. 2 c) inset, the apparent overlap seems to be smaller compared to that of Fig. 2 a) or b). However, relative to the minimum domain (which is the teal colored non-kink fraction in Fig. 2c) inset), the overlap is quite significant. To circumvent this issue, we have decided to compute a overlap metric while taking only the minimum domain in consideration. Hence the modified equation 2 becomes the following:

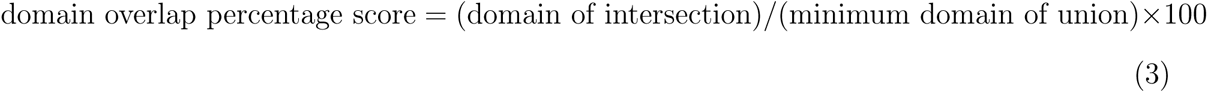

Despite the varying ways of quantifying the overlaps we have observed that *θ_B_* vs. *θ_R_* combination provides the best overlap score in all three terms when compared to pairwise combinations of other structural parameters as seen in Fig.2 d).

### Sequence filter

Our sequence filter is a literature-curated set up. For a 4 residue window, the central residues are either Glycine or aromatic amino acids. The terminal residues can be Glycine or aromatic amino acids or polar amino acids like Serine, Asparagine, Glutamine. The kinks are enables for neighbouring aromatic amino acids to form *π*-*π* interactions within adjacent stacks.

### Metadynamics simulation of wild type and mutant ^259^*SY NDFG*^264^ motif

The structure wild type ^259^*SY NDFG*^264^ motif obtained from one of the PrLD conformation containing a *β*-kink motif. We performed the *D*262*V* mutation using Pymol to obtain the mutant conformation.^47^ Both of the systems were then solvated with TIP4Pd water molecules and neutralized using a salt concentration of 150 mM NaCl. We have used a99SB-disp for equilibration and simulation.^48^ Both the systems were energy minimized and equilibrated to a temperature of 310 K and 1 atm pressure. As for production run, we have used the velocity rescaling algorithm to maintain the temperature.^49^ We have used LINCS algorithm to generate h-bond constraints.^50^ For the long-range electrostatic interactions a particle mesh Ewald scheme was used with order 4 and a Fourier spacing of 0.16. As for the short-range interactions, Verlet scheme was used with a cutoff of 1.2 nm. Both the simulations were performed using GROMACS-2022.5^51^ with PLUMED-2.9.1 patch.^52^ The details of both the systems and respective convergence plots can be found in the Supplemental Information (SI fig.10 a),b)). Please note that we have used *θ_B_* of ^260^*Y NDF* ^263^ window as our collective variable for the meta-dynamics simulation.

## Data availability

The full trajectory data of RGG and PrLD simulations considered in this work are available on our server for download. The SVM model and relevant codes are also available on our server. The server data can be publicly accessed via our laboratory GitHub link: codesrivastavalab/. The files can also be accessed directly from the publicly available Figshare portal: https://figshare.com/s/ea0cfee10359936eb3e0

## Acknowledgments

The High-Performance Computing facility “Beagle” setup from grants by a partnership between the Department of Biotechnology of India, the NSM supercomputer of IISC-Parampravega, Canada’s national high performance computing system Compute Canada are greatly acknowledged. AS thanks the DST for the National Supercomputing Mission grants (DST/NSM/R&D-HPC-Applications/2021/03.10, DST/NSM/R&D-HPC-Applications/Extension Grant/2023/27). AS would also like to thank the Teams Science Grant from the DBT-Wellcome Trust India Alliance (Grant number: IA/TSG/21/1/600245). AS also thanks the DBT National Network Project (NNP) grant on membrane-AMP interactions (BT/PR40323/BTIS/137/78/2023) and the Matrics grants on peripheral membrane protein (MTR/2023/001040) from the Science and Engineering Board (SERB), India.

IR and AS thank Prof. George Rose (Johns Hopkins University) for critically reading and commenting on the article. IR, RA and AS dedicate this work to late Prof. N. Srinivasan whose counsel we sorely missed during this work.

## Author contributions

A.S., I.R. and R.A. conceptualization; I.R. and A. S. methodology; I.R analysis; I.R. investigation; I.R. visualization; I.R. and A.S. writing–original draft; A.S., I.R. and R.A. writing–review & editing; A. S. supervision.

## Funding and additional information

A. S. thanks the Department of Science and Technology for the National Supercomputing Mission grant (Department of Science and Technology/NSM/RD high-performance computing [HPC]; Applications/2021/03.10) and the Supercomputing Education and Research Center at IISc-Bangalore for the HPC resources. I.R. acknowledges financial support from the NSM grant. A. S. and I.R. also acknowledge the access to the computing facility from the Compute Canada and the consumable grant from the IISc-IoE that supported the access to the facility.

## Supporting Infomation

### Visualization of training datasets

In Fig. 1 d), e) we have plotted different combinations of structural collective variables to visualise our training datasets. In both *C*1 *− C*4 vs *θ_B_* and *C*1 *− C*4 vs *θ_R_* (see SI Fig. 1 d),e)) the kink (blue) and non-kink(red) regions overlap into multiple ambiguous domains. However, the combination between *θ_B_*-*θ_R_* stands out by resolving the ambiguity through creating another bifurcating branch in the kink domain (see Fig. 2 b) highlighted by the red circled region). This is better understandable in a 3D plot of *C*1 *− C*4 vs *θ_B_* vs *θ_R_* of the training data sets where only along the *θ_B_*-*θ_R_* plane the bifurcation of the new branch of the kink from the ambiguous region is visible(see movie_1_*.mp*4).

To emphasize that each individual collective variables are insufficient for classification we have also provided the respective histograms in SI Fig. 1 a), b), c). Note that in SI Fig. 3 a), b), c) we have also plotted the *C − N − CA − CB* torsion angle histogram and *C − N − CA − CB* vs *θ_B_* and *θ_R_* separately in a 2D space. Although there is an apparent separation between the clusters of two motifs we should note that *C − N − CA − CB* torsion angle is not a generalised parameter that is well-defined for every residue. Because it considers side chains which are absent in Glycine. This is relevant as we have observed Glycine to be quite prevalent among the *β*-kink promoting residues and generally lies in the region of ambiguity which usually creates the bifurcating branch (highlighted by red circle) from the kink domain in the *θ_B_*-*θ_R_* space. This can be observed from the 3D plot in Fig. 2 a) where the absence of side chains in Glycine is causing the disappearance of the bifurcating branch (because it is undefined) which is clearly visible in the 2D plot of *θ_B_* vs *θ_R_* Fig. 2 b).

This establishes that none of the parameters just by itself does provide a confident classification between the kink and non-kink conformations (as explained in the histograms in Fig. 1 a), b), c) and Fig. 3 a). Also the pair-wise combination of all these parameters (except *θ_B_*-*θ_R_*) (Fig 1 d), e) and Fig. 3 a), c), d)) are unable to resolve the ambiguous regions in any further subdomains. Even after the successful segregation of regions in *θ_B_*-*θ_R_* space, still some ambiguous points remain near the decision boundary. We speculate that the time evolution of any of these structure within the ambiguous region may lead to either a stable kink or a non-kink point.

Another important point to be noted is the distribution of *ψ*-*ϕ* angles from kink and non-kink data set in the Ramachandran map. We can see that the kinked data points are all over the Ramchandran map. Some of the points are present even in the standard *β*-sheet region. This could be due to the influence of side chain or the lack thereof. As a result even though the *ψ*-*ϕ* angles are falling in the known *β*-sheet region, the side-wise tilt of the back bone could be giving rise to the kinks. Additionally, from the training data set as well as from the test data set six clear clusters of kinks are visible in the Ramachandran map.

**Fig. S 1:**
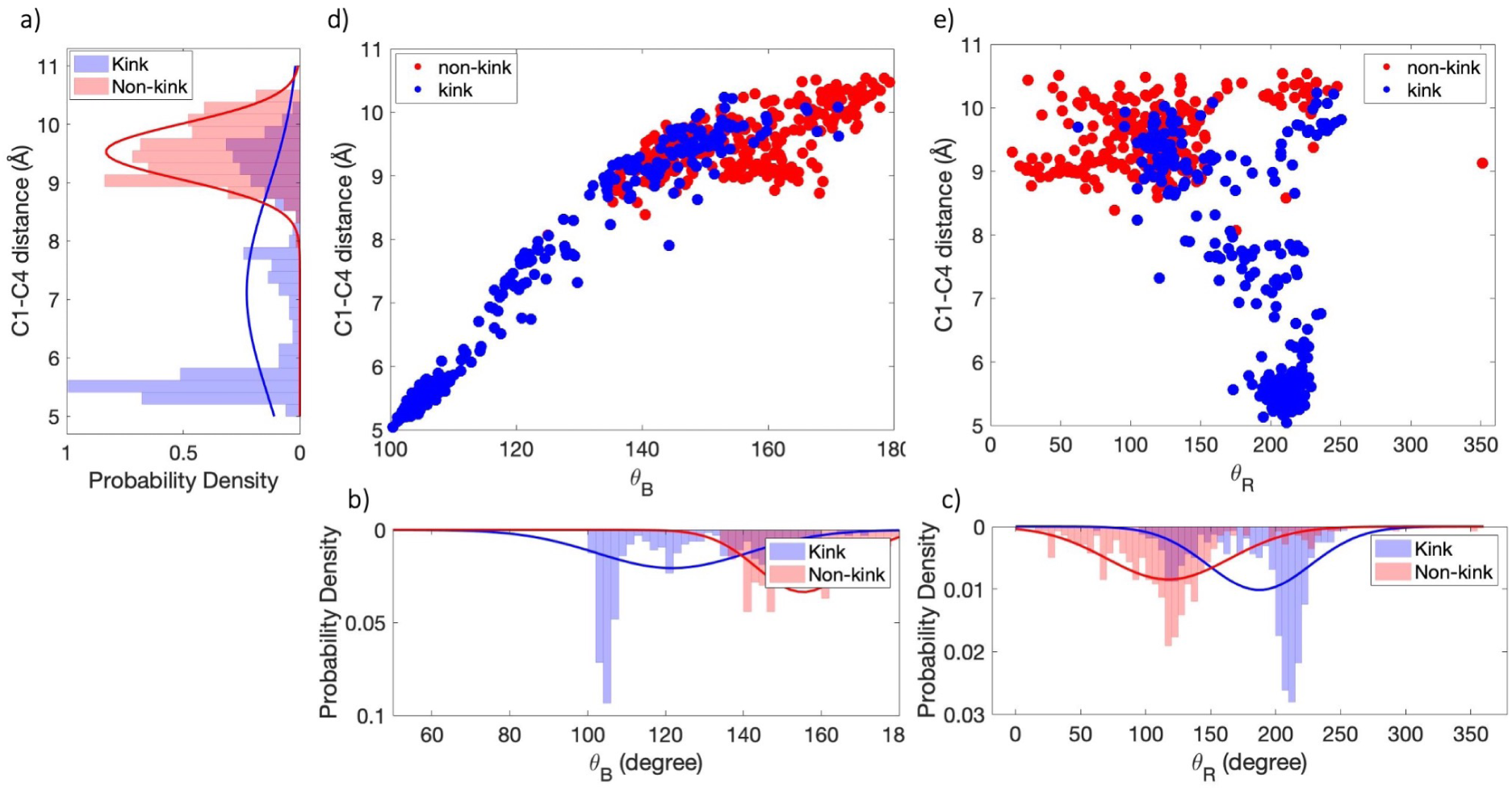
The probability distribution of kink(blue) and non-kink(red) dataset with respect to different collective variables. a) Probability density of *C*1 *− C*4 distance b) Probability density of *θ_B_* c) Probability density of *θ_R_*. The phase space arrangement of data points from training data sets with blue markers being kinks and red markers being non-kinks. d) *C*1 *− C*4 distance vs *θ_B_* e) *C*1 *− C*4 distance vs *θ_R_*

**Fig. S 2:**
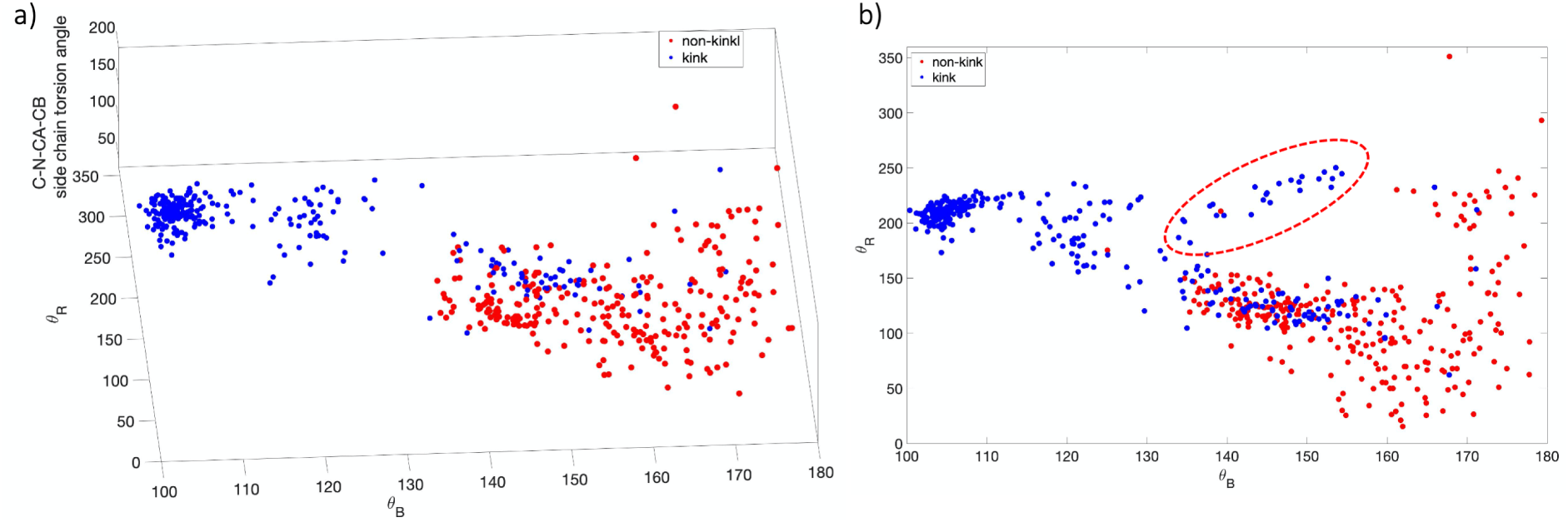
The phase space arrangement of data points from training data sets with blue markers being kinks and red markers being non-kinks. a) *C − N − CA − CB* distance vs *θ_B_* vs *θ_R_* b) *θ_B_* vs *θ_R_*

**Fig. S 3:**
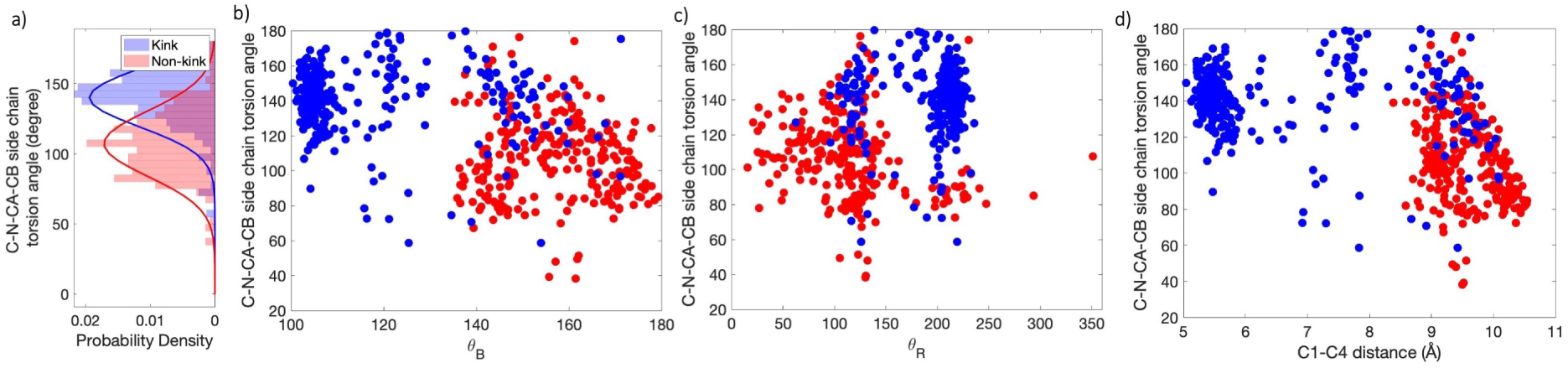
a) The probability distribution of kink(blue) and non-kink(red) dataset with respect to *C − N − CA − CB* torsion angle. The phase space arrangement of data points from training data sets with blue markers being kinks and red markers being non-kinks. d) *C − N − CA − CB* distance vs *θ_B_* e) *C − N − CA − CB* distance vs *θ_R_*

**Fig. S 4:**
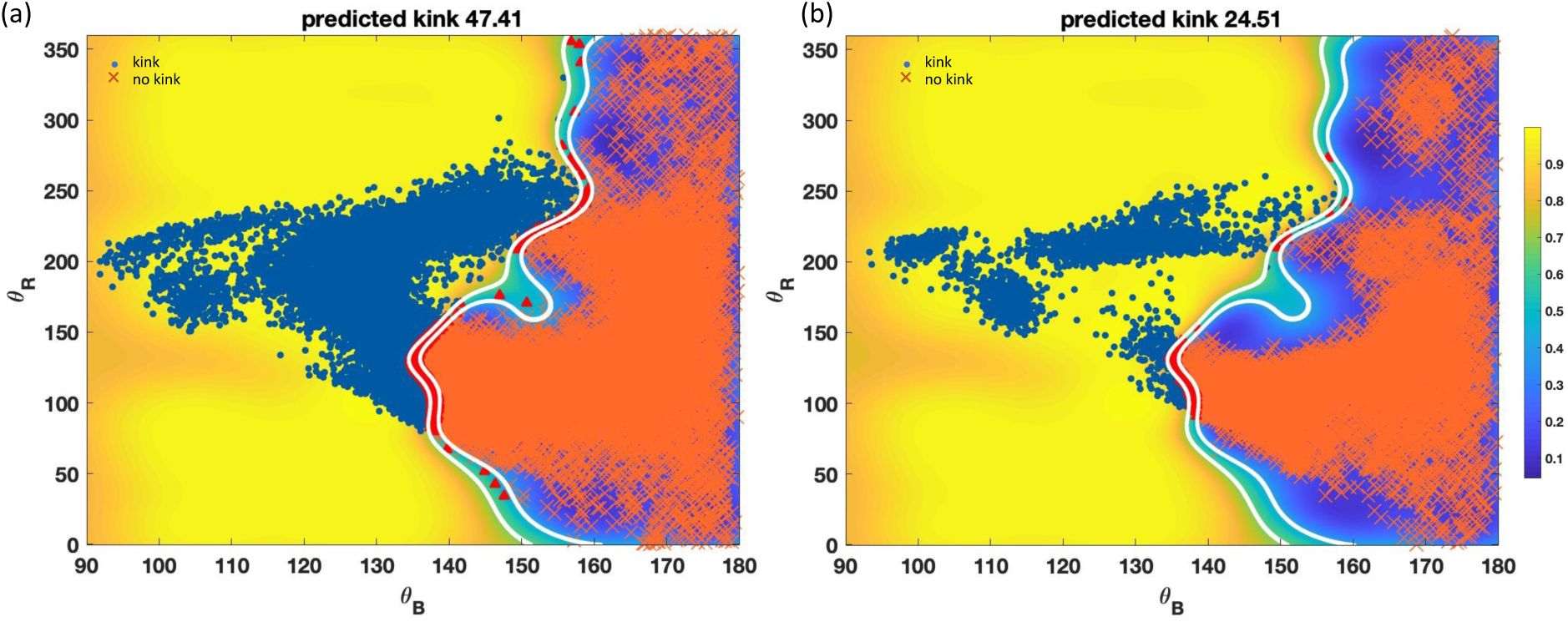
Kinked-*β* strands classified based on *θ_B_* and *θ_R_* angles using SVM for simulation datasets. a) *α*-synuclein (1-140), ^1^ b) Amyloid-*β* (1-42).^2^ The color map indicates the probability of kink or non-kink spanning from yellow to blue (kink to non-kink) with boundary at 0.5 (green). The blue markers stand for kink and the orange markers are for non-kink.

**Fig. S 5:**
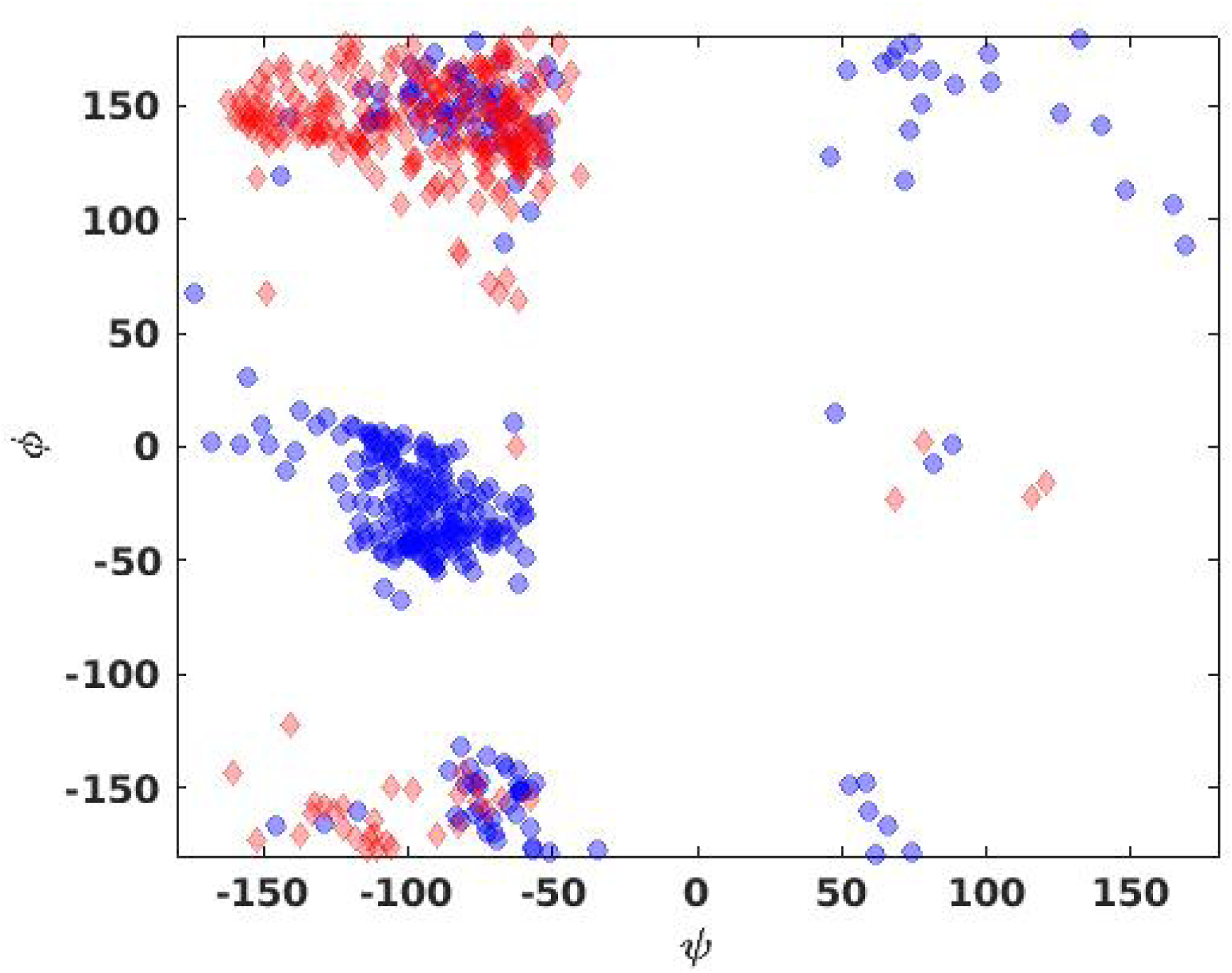
The *ψ*-*ϕ* angles distribution of kink(blue) and non-kink(red) data set from training data.

**Fig. S 6:**
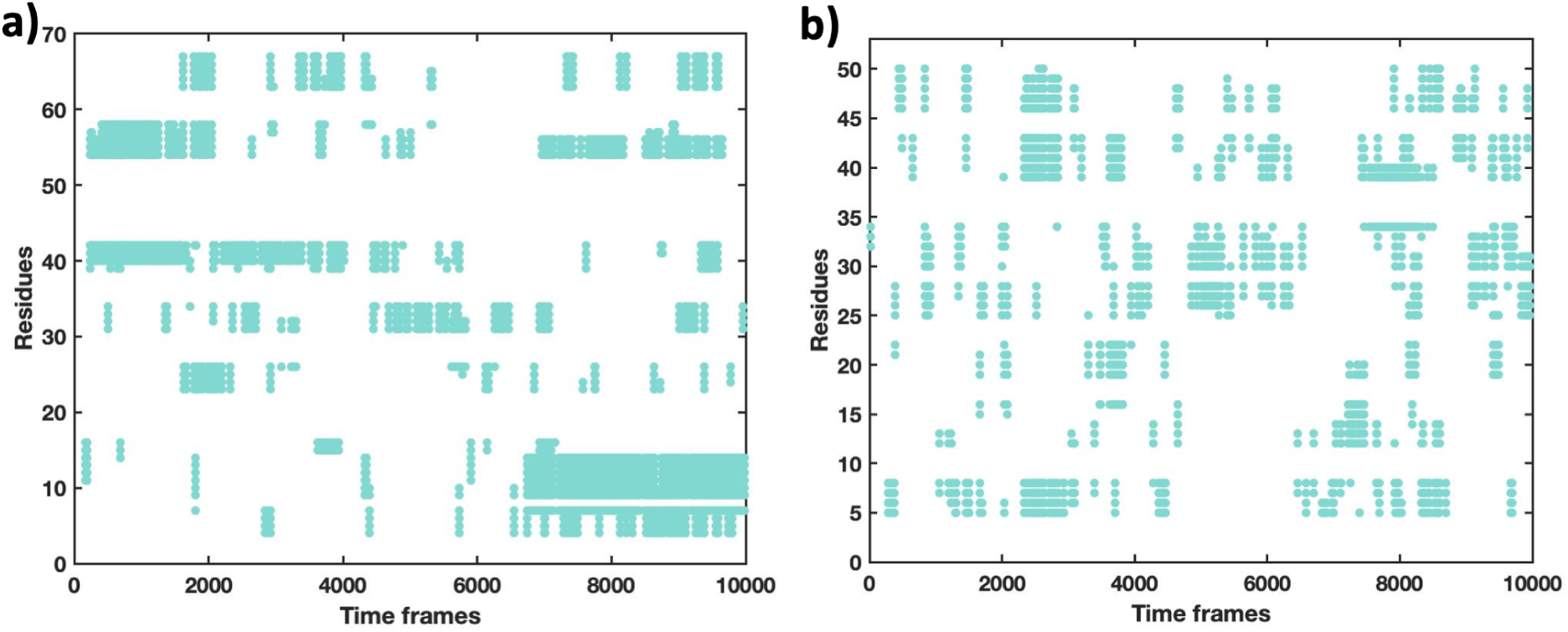
LARKS profile plot with respect to time evolution for a) PrLD and b) RGG. On X-axis and Y axis, time and residues are present, respectively.The markers flag the LARKS forming residues across time.

**Fig. S 7:**
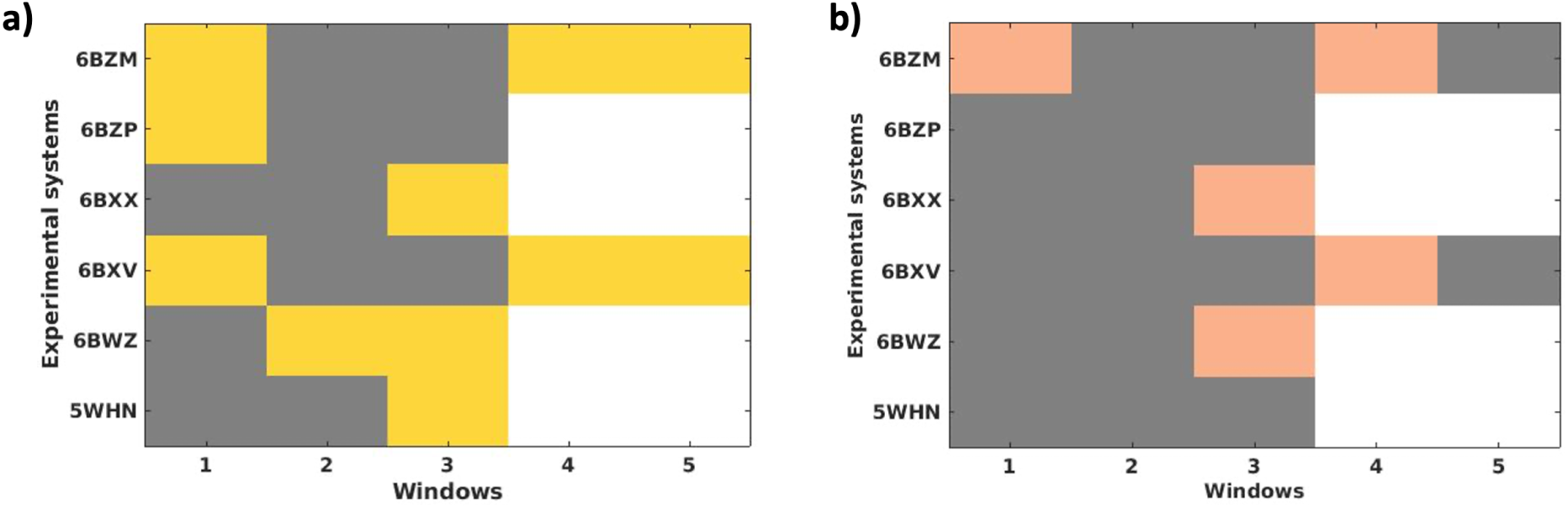
a) This is a derivation of the MS fig.6 d) which shows presence or absence of kinks for the 6 experimentally solved crystal structures. The yellow (grey) blocks for each residue window indicate presence (absence) of kink. Note that, all the experimental systems are not of equal in length. Hence the systems with white blocks at the edge indicate end of the windows. b) Presence and absence of LARKS in experimental data. The peach (grey) blocks for each residue window indicate presence (absence) of kink. Note that, all the experimental systems are not of equal in length. Hence the systems with white blocks at the edge indicate end of the windows. The sequence filter used here is mentioned in MS Table - 1.

**Fig. S 8:**
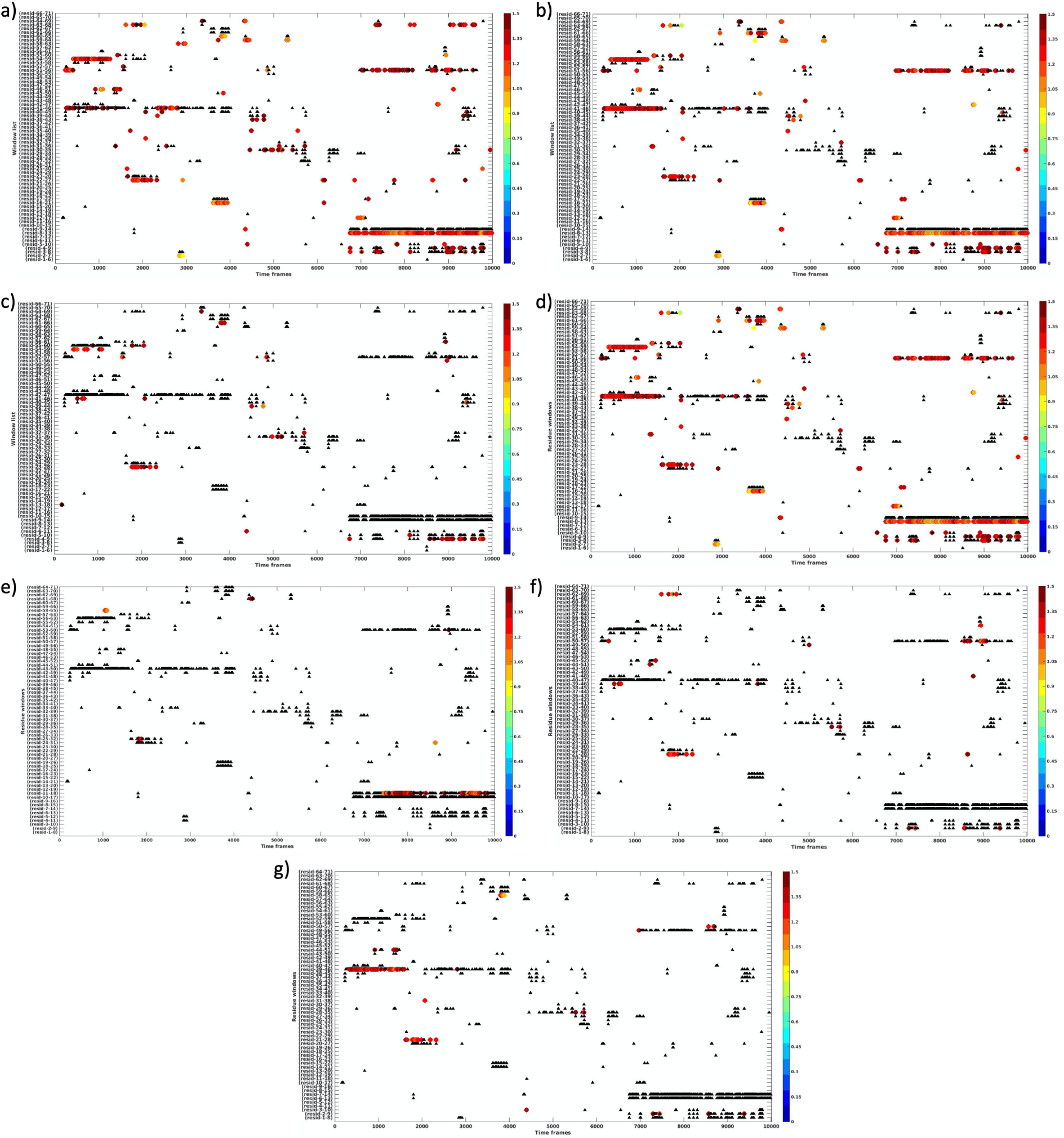
Peptide segments profile for PrLD within 1.5Å RMSD cutoff while taking experimental data as reference. a) 5WHN: residue-window 3, b) 6BWZ: residue-window 2, c) 6BWZ: residue-window 3, d) 6BXX: residue-window 3, e) 6BXV: residue-window 1, f) 6BXV: residue-window 4 g) 6BXV: residue-window 5. The black triangles are the population of all the segments having kinks at the same position with that of the experiment mentioned above. The residue window number indicates the particular residue window in the respective experimental data which produces a pair of *θ_B_*-*θ_R_* angle agreeing to the definition on kink. The coloured markers are the section the kinked-peptide segments within the RMSD value of 1.5Å. The color map stands for RMSD values ranging from 0-1.5Å with blue being the lowest and red being the highest value.

### RMSD comparison within simulation and experimental structures and creation of peptide library

We have performed the comparison between PrLD and RGG simulation data and each experimentally solved crystal structures to extract the conformations within 1.5Åcutoff. Among six available experimental structures we are reporting four of these structures for the comparison purpose. The hits of peptide segments profile with the threshold are plotted for both PrLD and RGG data with respect to each of the five experimental systems (SI Fig. 8 (for PrLD) and SI Fig. 9 (for RGG)).

**Fig. S 9:**
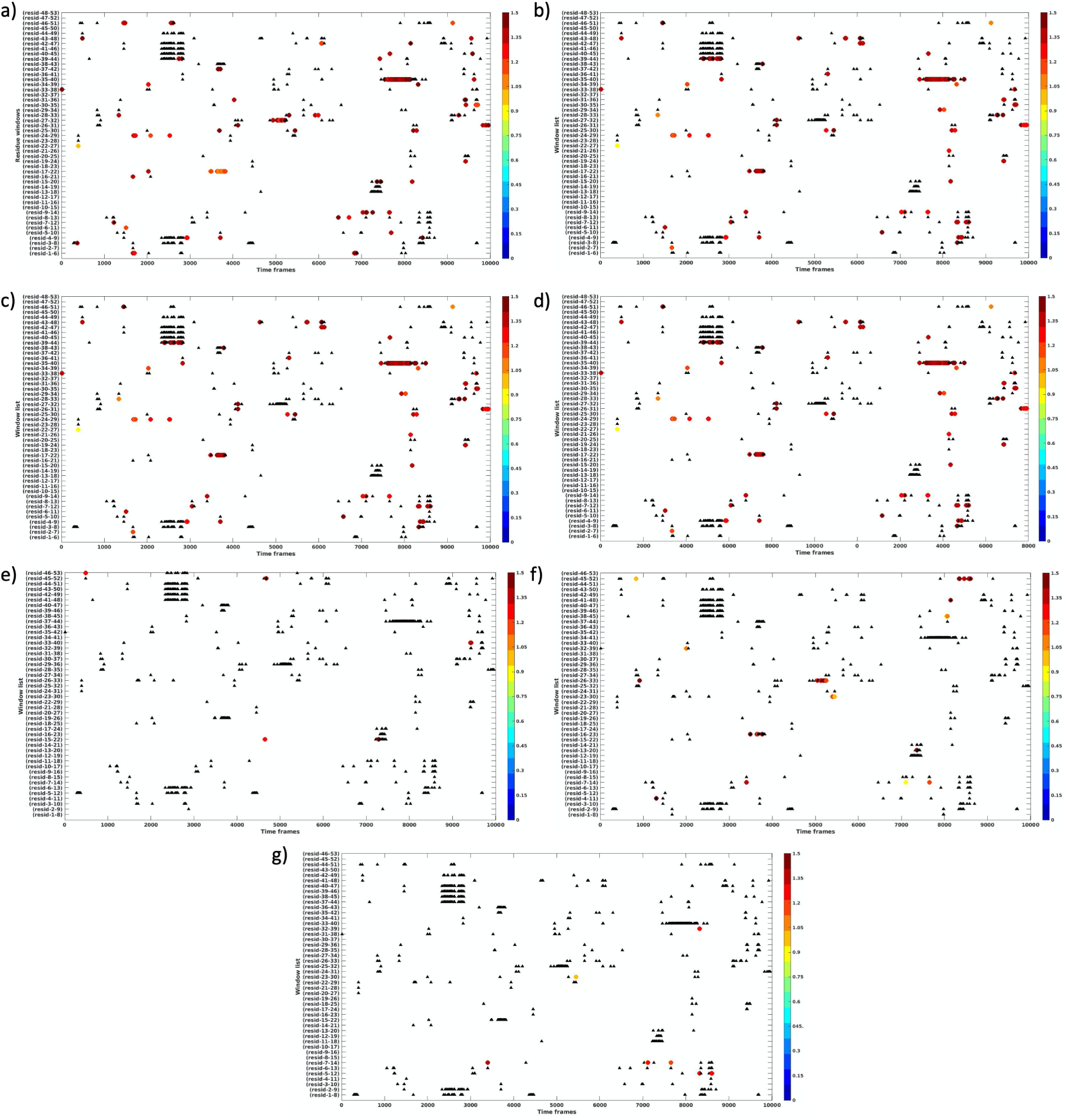
Peptide segments profile for RGG within 1.5Å RMSD cutoff while taking experimental data as reference. a) 5WHN: residue-window 3, b) 6BWZ: residue-window 2, c) 6BWZ: residue-window 3, d) 6BXX: residue-window 3, e) 6BXV: residue-window 1, f) 6BXV: residue-window 4 g) 6BXV: residue-window 5. The black triangles are the population of all the segments having kinks at the same position with that of the experiment mentioned above. The residue window number indicates the particular residue window in the respective experimental data which produces a pair of *θ_B_*-*θ_R_* angle agreeing to the definition on kink. The coloured markers are the section the kinked-peptide segments within the RMSD value of 1.5Å. The color map stands for RMSD values ranging from 0-1.5Å with blue being the lowest and red being the highest value.

**Fig. S 10:**
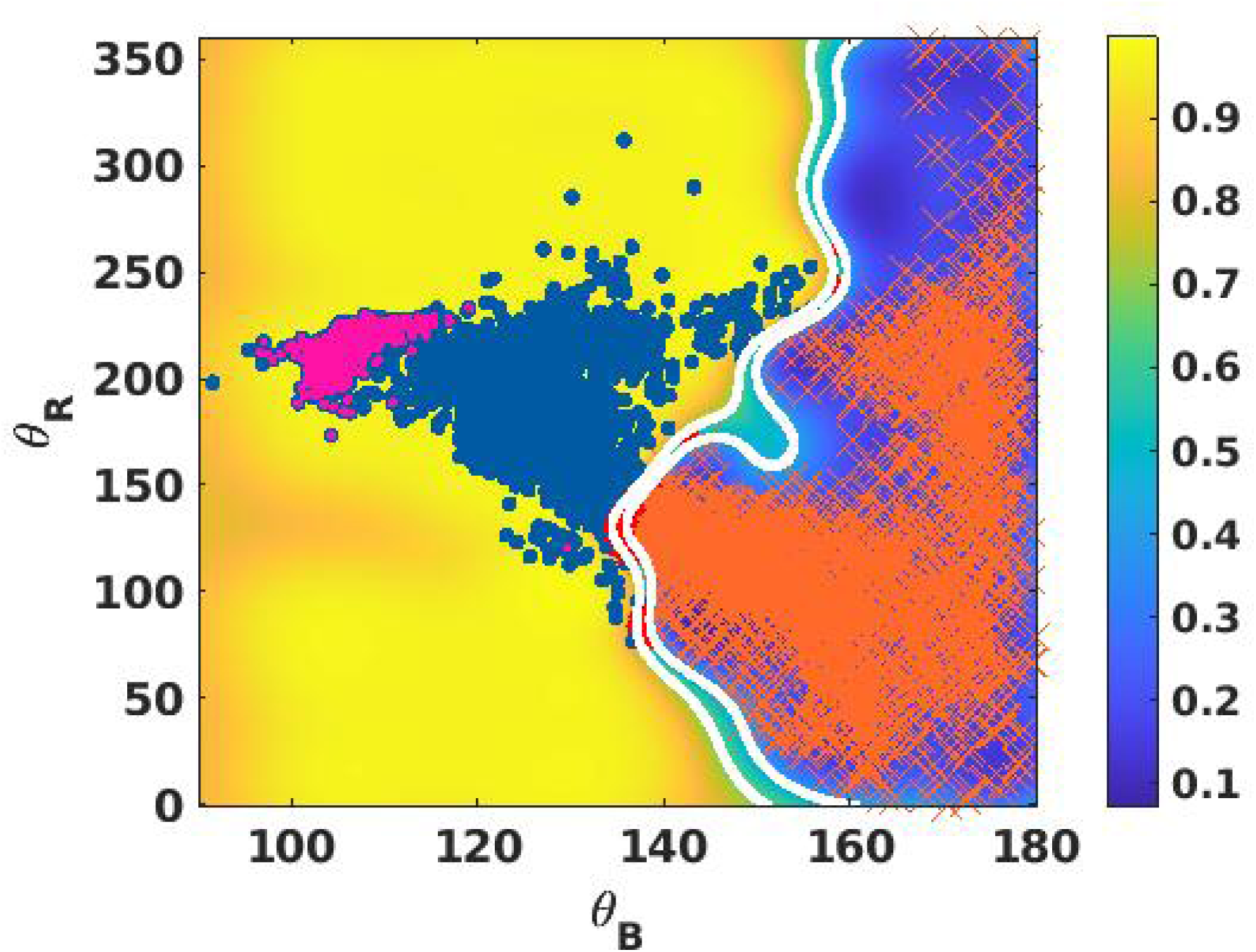
Data points containing *β*-turn like features are shown as pink markers in the *θ_B_*-*θ_R_* SVM map. Yellow space is the kink region where blue markers are kink data points. Blue space is non-kink region where orange marker stand for non-kink data. Green region is decision boundary which is outlined by two white lines indicating the ambiguous region.

**Table 1:** System information of wild type and mutant ^259^SYNDFG^264^ segment used for metadynamics.

| Peptide sequence | System size | Time(ns) |
| --- | --- | --- |
| $^{259}\text{SYNDFG}^{264}$ | 6041 | 300 |
| $^{259}\text{SYNVFG}^{264}$ | 6854 | 300 |

**Fig. S 11:**
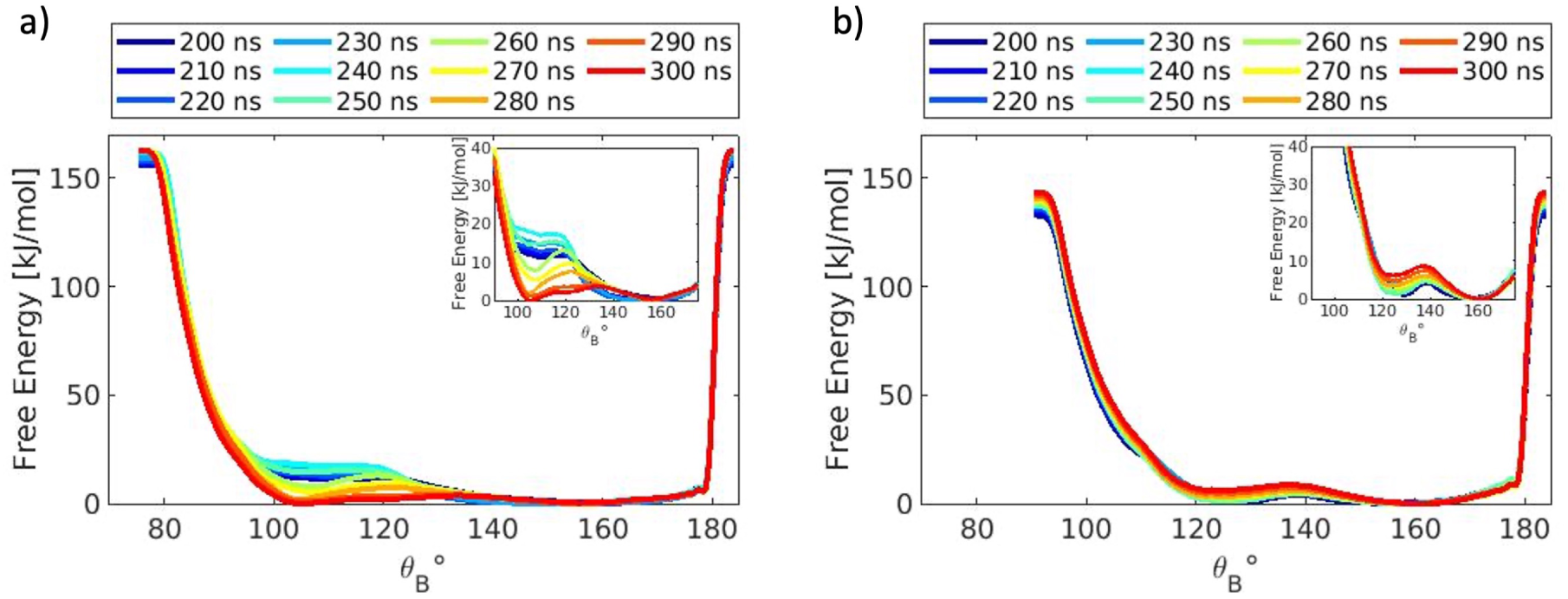
Free energy vs. *θ_B_* plot showing convergence for metadynamics simulation of a) wild type ^259^*SY NDFG*^264^ peptide segment and b) mutant ^259^*SY NV FG*^264^ peptide segment. On the X-axis *θ_B_* is plotted. On the Y-axis free energy is plotted. These plots are made from last 100 ns of the simulation. The time stamps are color code from blue to red in an ascending manner. A fraction of both the plots are zoomed in as insets to emphasize the difference between the wild type and mutation data.

